# Structural organization of p62 filaments and the cellular ultrastructure of calcium-rich p62-enwrapped lipid droplet cargo

**DOI:** 10.1101/2024.10.15.618463

**Authors:** Sabrina Berkamp, Lisa jungbluth, Alexandros Katranidis, Siavash Mostafavi, Olivera Korculanin, Peng-Han Lu, Lokesh Sharma, Lipi Thukral, Jörg Fitter, Rafal E. Dunin-Borkowski, Carsten Sachse

**Author notes:** Correspondence (C.S.). These authors contributed equally (Sabrina Berkamp, Lisa jungbluth, Alexandros Katranidis).

## Abstract

The selective autophagy receptor p62/SQSTM1 (from hereon p62) is known to form higher-order filaments in vitro and to undergo liquid-liquid phase separation when mixed with poly-ubiquitin. We determined the full-length cryo-EM structure of p62 and elucidated a structured double helical filament scaffold composed of the PB1-domain associated with the flexible C-terminal part residing in the lumen and the solvent-accessible major groove. At different pH values and upon binding to soluble LC3, LC3-conjugated membranes and poly-ubiquitin, we observed p62 filament re-arrangements in the form of structural unwinding, disassembly, lateral association and bundling, respectively. In the cellular environment, under conditions of ATG5 knockdown leading to stalled autophagy, we imaged high-contrast layers consisting of p62 oligomers enwrapping lipid droplets by cryogenic electron tomography in cellulo, which we identified as calcium phosphate by compositional spectroscopy analysis. Together, we visualized the cellular ultrastructure of p62 oligomers with high calcium content as a potential intermediate step of autophagy initiation.

## Introduction

Autophagy is a conserved cellular process that degrades and recycles cytoplasmic material such as aggregated proteins, damaged organelles and pathogens to maintain intracellular homeostasis (Mizushima et al., 2011; Wen & Klionsky, 2016). While autophagy operates at a basal level for routine maintenance, it can also be activated by stress factors like nutrient deprivation, hypoxia and DNA damage (Kroemer et al., 2010). There are three main autophagy pathways: macroautophagy, microautophagy and chaperone-mediated autophagy. In macroautophagy (from here on simply referred to as autophagy), an autophagosome forms within the cytoplasm to enclose cellular material that is subsequently degraded and recycled after fusing with a lysosome. A series of defined autophagy core machinery complexes mediate the progression of cellular autophagy: Atg1/ULK1 kinase, PI-3 kinase, ATG9 lipid scramblase, the ATG2-ATG18 lipid transfer complex and two ubiquitin-like conjugation systems (Xie & Klionsky, 2007). The recognition of cargo can proceed in a non-selective or in a selective manner by either enclosing the bulk stochastically or through specific binding by selective autophagy receptors, respectively (Y. Feng et al., 2014).

Selective autophagy requires cargo receptors that recognize poly-ubiquitinated cargo and interact with Atg8 family proteins (Kraft et al., 2014; Y.-K. Lee & Lee, 2016). The Atg8 family contains ubiquitin-like proteins: Atg8 in yeast and microtubule-associated protein 1 light chain 3 (LC3) as well as gamma-aminobutyric acid type A receptor-associated protein (GABARAP) in mammals (Shpilka et al., 2011). The conjugation of Atg8 proteins to the lipid membrane is essential for phagophore membrane expansion (Mizushima et al., 2001) and is mediated by two autophagy-specific ubiquitin-like conjugation systems: ATG12-ATG5 and the LC3 conjugation system (Nakatogawa et al., 2009). Selective autophagy receptors, like p62/SQSTM1 (from here on: p62) (Bjørkøy et al., 2005), contain an LC3-interacting region (LIR) motif that facilitates interaction with Atg8 proteins and a ubiquitin-binding domain (Birgisdottir et al., 2013; Rogov et al., 2014). In the meantime, a series of additional autophagy receptors have been identified with LIR motifs as well as ubiquitin binding abilities in order to confer specificity and redundancy in the cargo recognition and removal. For instance, for mitophagy NBR1, optineurin, NDP52 and TAXBP1 have been characterized in addition to p62 (Lazarou et al., 2015). While p62 knock-outs are viable for an organism, long-term benefits of increased p62 levels have been found to promote proteastasis and longevity in response to stress in C*.elegans* (Kumsta et al., 2019).

p62 has a key role in selective autophagy by recognizing poly-ubiquitinated cargo and delivering it to the autophagosome (Bjørkøy et al., 2005). Originally, p62 was described as an interaction partner for tyrosine-protein kinase Lck (Joung et al., 1996), while it was also shown to be involved in multiple pathways including Wnt, Nrf2, mTORC and NF-κB signaling (Puissant et al., 2012). In the NF-κB pathway, p62 can interact with MEKK3, MEK5, aPKCs (Lamark et al., 2003) and tumor necrosis factor receptor associated factor 6 (TRAF6) as a mediator in tumorigenesis (Duran et al., 2011). Moreover, p62 actively promotes the spatio-temporal organization of autophagy downstream effectors such as FIP200 at the cargo site (Turco et al., 2019). p62 is a multi-domain protein with a Phox1 and Bem1p (PB1) domain, followed by a ZZ-domain and an intrinsically disordered region (IDR) up to the C-terminal ubiquitin-binding associated (UBA) domain. The PB1 domain is responsible for p62 polymerization into long flexible filaments (Ciuffa et al., 2015; Jakobi et al., 2020) and was shown to be critical for its function in autophagy as polymerization deficient mutations excluded p62 from the autophagosome formation site (Itakura & Mizushima, 2011). The IDR contains several relevant interaction motifs, including the LIR and KEAP1-interacting region (KIR) motif (Komatsu et al., 2010). The C-terminal UBA domain captures poly-ubiquitinated cargo (Ciani et al., 2003). The autophagy receptor NBR1 is structurally the most similar to p62 amongst the autophagy receptors as it is closely evolutionarily related and directly interacts with p62 via its PB1 domain albeit it is unable to form higher-order oligomers or filaments (Jakobi et al., 2020; Kirkin et al., 2009).

p62 is a well-studied cargo receptor that recognizes and removes various different cellular cargoes: aggregated proteins in aggrephagy (Bjørkøy et al., 2005; Pankiv et al., 2007), damaged peroxisomes in pexophagy (Zhang et al., 2015) and lipid droplets in lipophagy (L. Wang et al., 2017; Yan et al., 2019). For lipophagy, the size of the removed lipid droplets is smaller than one µm while larger lipid droplets were shown to be predominantly degraded by lipolysis (Schott et al., 2019). At the same time, lipid droplets are a special cargo in the context of autophagy as they can also serve as a direct or indirect lipid source for the growing phagophore mediated by direct contact in addition to the ATG9A vesicles or ATG2 lipid bridges (Dupont et al., 2014; Korfhage et al., 2023; Mailler et al., 2021; Shpilka et al., 2015). Lipid droplets were also found to serve as direct substrates for non-canonical autophagy during prolonged starvation after LC3-conjugation (Omrane et al., 2023).

p62 has also been shown to undergo liquid-liquid phase separation (Brangwynne et al., 2009) induced by poly-ubiquitin interaction with multiple polymerized p62 entities in vitro (Sun et al., 2018; Zaffagnini et al., 2018a). However, in vivo, p62 phase separated condensates recruit many autophagy regulators that can affect the mobility of the separated phase and as a result autophagy efficiency. For instance, Keap1 over-expression was observed to increase the rigidity of p62 condensates, thus promoting autophagy (Y. Lee et al., 2017). NBR1 enhances p62 condensate formation and poly-ubiquitin affinity (Turco et al., 2021). In addition to binding partners, the oligomeric state of p62 and post-translational modifications can affect the phase separation properties, e.g., as the IDR of p62 harbors the KIR motif that when post-translationally modified has been shown to enhance the interaction with Keap1 (Ichimura et al., 2013). The quaternary structural organization of p62 presents another level of regulation for cellular phase separation as phosphorylation of the PB1 domain by PKA was identified and shown to disrupt PB1 domain interactions (Christian et al., 2014).

In order to improve our detailed structural understanding of purified p62, we determined the cryo-electron microscopy (cryo-EM) structure of full-length p62. The structure revealed the double helical organization of an N-terminal PB1-domain scaffold while the more poorly resolved partially disordered C-terminal part resided in the lumen and the major groove of the assembly. Binding studies of LC3b and poly-ubiquitin showed that p62 filaments can undergo structural re-arrangements of disassembly or filament bundling, respectively. When p62 filaments were added to LC3b-conjugated membranes, they associate laterally and also unwind their regular helical structures into bubbles at physiological pH. Native ultrastructural imaging of p62-positive structures using cryo-electron tomography (cryo-ET) in ATG5 depleted RPE1 cells revealed stalled intermediate autophagy stages of p62-enwrapped lipid droplets. Interestingly, the surrounding p62 layer had a dark contrast and contained high levels of calcium and phosphorus based on energy-dispersive X-ray spectroscopy analysis (EDX). The high-resolution visualization of p62-enwrapped cargo together with calcium may present an important intermediate state of autophagy initiation.

## Results

### p62 forms a double-helical filament with a PB1 scaffold and a flexible C-terminus in the major groove

Previously, we determined the three-dimensional (3D) cryo-EM structure of the p62-PB1 domain arranged in a series of different filamentous assemblies (Jakobi et al., 2020). In order to advance our structural understanding of full-length human p62, we optimized the previous preparations of full-length p62 for improved homogeneity. As previous studies suggested an effect of pH on the assembly conditions (Kwon et al., 2018), we compared the preparations at different pHs. Classifications of small cryo-

EM test data sets revealed that at pH=6 the majority of the filament classes were helically regular while at pH=8 a large fraction of them were found unwound resembling an open bubble (**Supplementary Fig. 1A**). Therefore, we recorded a full cryo-EM data set at pH=6 revealing long and flexible filaments with a width of approximately 15 nm (**Figure 1A**). Using segmented helical reconstruction (Desfosses et al., 2014; Punjani et al., 2017; Scheres, 2012), we set out to determine the structure of p62 filaments. Due to pitch heterogeneity between 130 and 160 Å (**Figure 1B** and **Supplementary Fig. 1B**), we refined a more homogeneous subset with a 150 Å pitch and 13.6 units per turn (corresponding to 11.0 Å helical rise and a −26.1° helical rotation) including dihedral symmetry and obtained a 3D reconstruction at 4.5 Å global resolution (**Supplementary Fig. 1C-D, Table 1**). Consistent with the class averages, the local resolution varied significantly from a well-defined PB1 domain scaffold at 4.2 Å resolution up to over 9 Å for the more flexible regions (**Figure 1C**). The cryo-EM structure revealed a left-handed double-helical filament with two antiparallel strands of PB1 domains in the full-length p62 assembly. The atomic model of the PB1 domain of p62 (PDB: 6TGY) fitted well in the best defined scaffold density supporting β-strand separation (**Figure 1D left**). At lower density thresholds, the ZZ domain envelope emerged connected to the PB1 domain albeit at lower resolution (**Figure 1D right**). The densities of the associated C-terminal domains were weak and poorly resolved presumably due to flexibility with respect to the PB1 strands. Based on our p62 filament structure, we expanded the domains to match the PB1 density to the experimentally observed helical symmetry assembly (**Figure 1E**). The resulting p62 double helix architecture is built on the scaffold of the PB1 domain generating a major groove that can accommodate the flexible C-terminal part.

**Figure 1:**
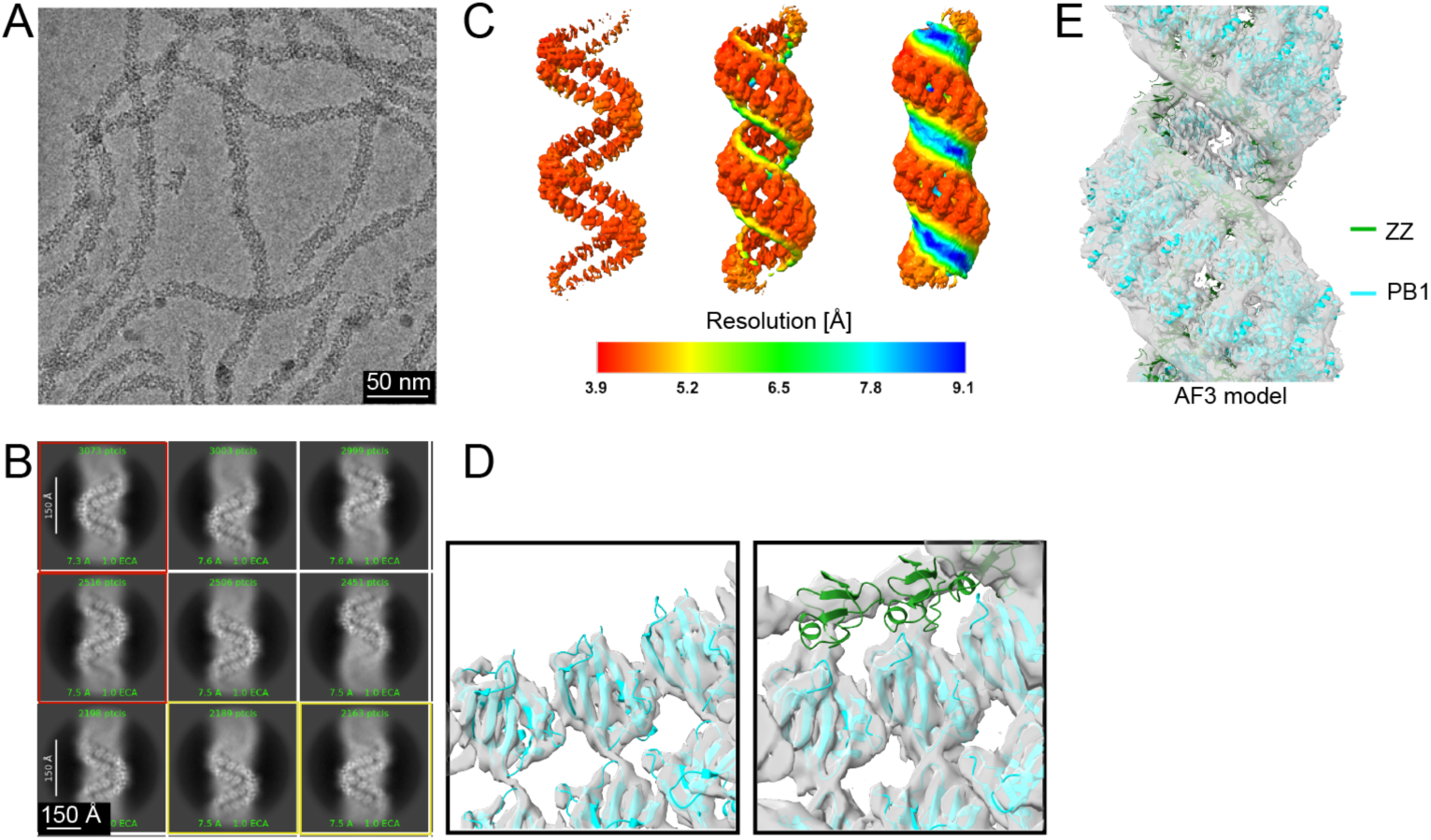
Cryo-EM structure and atomic model of full-length p62 filaments. (A) Representative cryo-micrograph with long and flexible p62 filaments. (B) 2D classes showing well-defined densities of the filamentous scaffold in addition to blur including apparent variation in pitch, with higher helical pitch classes (red frame) and lower helical pitch classes (yellow frame). (C) Three-dimensional density of p62 superimposed with local resolution values showing PB1 domains arranged in a left-handed double helical architecture. The map rendered at different thresholds shows that the PB1 domain is well-resolved (∼4.5 Å resolution), the ZZ domain appears as a featureless blob at ∼6.5 Å resolution and the remaining C-terminal domains are not well resolved. (D) Left: The atomic model of the PB1 domain fits well into well-defined PB1 density at higher threshold. Right: The atomic model of the ZZ domain of p62 fits in the featureless blobs seen at lower threshold. (E) The atomic model of the helical p62 filament with highlighted PB1 (cyan) and ZZ (green) domains fitted into the filament density displayed at low threshold.

**Table 1:** Cryo-EM structure determination of full-length p62.

|  |  |
| --- | --- |
| Processing type | Helical segments |
| Microscope | Talos Arctica 200 keV |
| Detector | Gatan K3 |
| Magnification | 100,000x |
| Defocus range [μm] | -0.5 to -3.0 |
| Physical pixel size [Å] | 1.6778 |
| Total dose [e<sup>-</sup>/Å<sup>2</sup>] | 50 |
| Micrographs | 4038 |
| Extraction box size [Å] | 503 |
| Overlap [%] | 90 |
| Initial no. of segments | 1,000,000 |
| Final no. of segments | 600,000 |
| Helical symmetry |  |
| Pitch [Å] | 150 |
| Rise [Å] | 10.985 |
| Rotation [°] | -26.141 |
| Global map resolution (FSC=0.143)/FDR-FSC [Å] | 4.5/4.5 |

### p62 filament interactions with LC3b and ubiquitin

To experimentally study the interactions of p62 filaments with LC3b and ubiquitin, we employed single-particle fluorescence microscopy in total internal reflection fluorescence (TIRF) mode and negative staining EM. We formed p62 filaments at pH=8 and subsequently labeled them at lysine residues with multiple Cy5 dyes (p62 filaments-Cy5) while cyan fluorescent protein (CFP) was fused to LC3b (CFP-LC3b). Consistent with the observed p62 filaments, elongated p62 structures were visible in the Cy5 channel, rather mobile and diffusing above the surface in dilute nM concentrations on the cover slip (**Figure 2A** and **Supplementary Video 1**). Upon addition of equimolar amounts of CFP-LC3b, the same filament became visible in the CFP channel as well (**Figure 2B** and **Supplementary Video 2**) supporting that LC3b was bound to the p62 filaments. Upon addition of GST-4xUbiquitin (GST-4xUb) to the labeled filaments, we observed a rapid and spontaneous formation of µm-sized condensates that displayed minimal mobility (**Figure 2C** and **Supplementary Video 3**). Interestingly, the subsequent addition of CFP-LC3b to the formed condensates, resulted in the appearance of smaller mobile diffusing fragments (**Figure 2D** and **Supplementary Video 4**).

**Figure 2:**
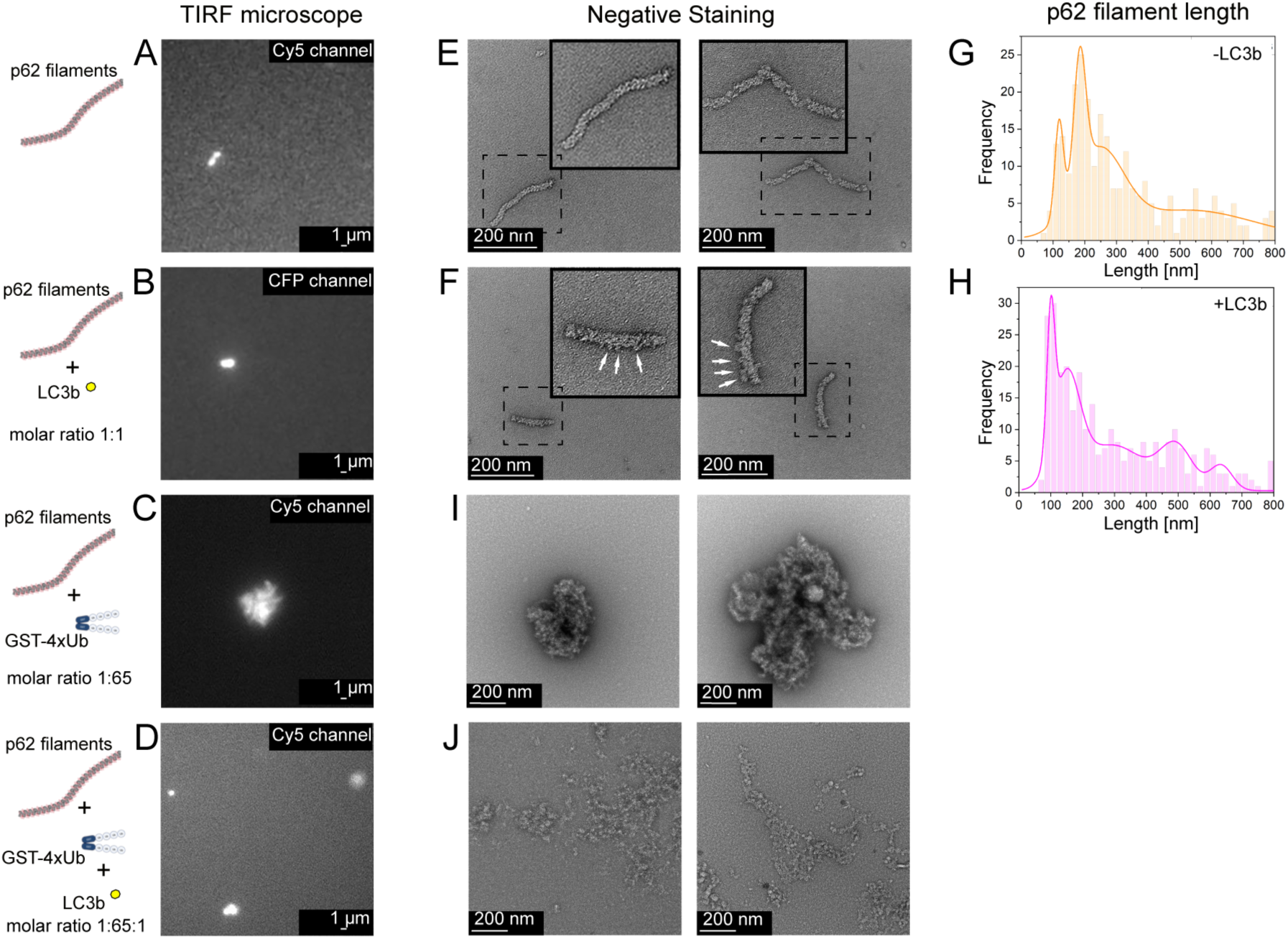
LC3b and poly-ubiquitin interactions with p62 filaments visualized by single-particle fluorescence and negative staining EM. p62 filament (A) labeled with Cy5 on TIRF microscope and (B) in the presence of CFP-LC3b. (C) p62 condensates forming in the presence of GST4x-ubiquitin (GST4x-Ub). (D) p62 condensates formed by GST4x-Ub disintegrate after addition of LC3b. (E) Negative staining of p62 filaments. Dashed boxes correspond to inset at higher magnification. (F) Negative staining of p62 filaments decorated with LC3b (sample as in B) (white arrows in inset). Histograms of (G) p62 filament length (H) after addition of LC3b. (I) Negative staining of p62 condensates (sample as in D with GST4x-Ub). (J) Negative staining of p62 condensates after addition of LC3b (sample as in D with GST4x-Ub).

The corresponding p62 filament samples in nM concentration were also imaged by negative staining EM. In the presence of CFP-LC3b, p62 filaments exhibited a distinctive decoration in comparison with the filaments alone (**Figure 2E/F**, white arrows in insets). A quantitative analysis of the filament length distribution revealed a shift towards shorter filaments for CFP-LC3b (**Figure 2G/H**), suggesting that interaction with LC3b induced depolymerization of p62 filaments. Similarly, in the presence of CFP-LC3b, GST-4xUb-induced condensates were disintegrated into smaller substructures (**Figure 2I/J**). In support of previous studies (Sun et al., 2019; Zaffagnini et al., 2018b), our single-particle studies in fluorescence and electron microscopy performed in diluted nM concentration showed that poly-ubiquitinated cargoes induce condensate formation while LC3b can promote disassembly of preformed p62 filaments as well as preformed condensates.

In order to molecularly assess the cryo-EM structure and the presented binding experiments, we generated a multimer AI-prediction of p62 using AlphaFold3 (AF3) (Abramson et al., 2024) consisting of ten subunits that formed closed rings mediated by the well-characterized PB1 domain interactions (**Supplementary Fig. 2A**). In an expanded model matching the helical symmetry, the C-terminal UBA domain fits tightly to the major groove next to the PB1 domain that is consistent with the cryo-EM density (**Supplementary Fig. 2B**). When the model is seen in top view, the PB1 domain scaffold and UBA domain are located at the outer rim of the filament while the ZZ domain is located on the inside. Notably, parts of the IDR, including the LIR and KIR binding motifs, are positioned towards the filament’s outside, supporting the observed molecular access of p62 interacting partners through the major groove of the filament. To assess the interaction accessibility within the p62 filament in more detail, we generated a second p62 multimer model in the above described way, this time with LC3b known to bind to the LIR motif of p62 (Pankiv et al., 2007). In the resulting model, LC3b is tightly fit in the filament’s major groove surface between the PB1 domain and covering the UBA domain (**Supplementary Fig. 2C**). The LIR of p62 is positioned outward and bound in the hydrophobic pocket 1 (HP1) and 2 (HP2) of LC3 in agreement with previously determined X-ray structures (Ichimura et al., 2008). A third p62 AF3-model with bound ubiquitin revealed that ubiquitin also resided in the major groove suggesting a competing mode between LC3b and ubiquitin binding. Together, the AF3-based integrative molecular model of the p62 filament supports the observed accessibility to binding partners of ubiquitin and LC3b while it also provides a molecular framework for the observed opposing higher-order structures of filament bundling and disassembly.

### p62 filament membrane interactions

In order to more closely mimic the cellular conditions, we performed the interaction studies at higher µM concentrations of p62 and investigated the binding of GST-4xUb and LC3b maleimide-conjugated to small unilamellar vesicles (SUVs). Subsequently, we visualized the natively preserved ultrastructures using cryo-ET. To better characterize the fine ultrastructure of p62 filaments on the image level, we recorded tomograms using the Volta phase plate for detailed high-contrast visualization. The resulting 3D tomograms showed 15 nm-wide flexible p62 filaments consistent with repeating pitches of 14 nm along the filament exhibiting regular major groove indentations matching the pH=6 conditions described above (**Figure 3A**). Closer inspection revealed overlapping filaments crossing underneath each other, branching filaments where individual PB1 scaffold strands are adopting a close to 90° bent as well as a cross-over reminiscent of Holliday junctions in double-stranded DNA. Due to the underfocus-induced white halo of phase plate surrounding the filament density, we turned back to conventional cryo-imaging at underfocus and closer to physiological pH 7.4 for the subsequent tomograms. Upon addition of GST-4xUb, p62 filaments were found to concentrate locally by forming bundles of parallel filaments supported by filament image segmentations (**Figure 3B**). Bundle formation was accompanied by the formation of µm-sized 3D agglomerates that were often too thick to be imaged by cryo-ET while visible by negative staining EM (**Supplementary Fig. 3**) being consistent with the previous in vitro observations of phase separated p62 bodies (Sun et al., 2019; Zaffagnini et al., 2018b). When we incubated p62 filaments with LC3b-conjugated SUVs, we observed unbound filaments as in the p62-only control in proximity to SUVs as well as SUV bound filaments (**Figure 3C**). When not bound, the p62 exhibited the typical filament appearance with a regular 14 nm pitch and major groove indentations as well as unwound stretches of the high pH=8 p62 sample alone (**Figure 3D**). When p62 filaments were found connected to the SUV density, they were either laterally associated along the membrane of the SUVs or bound through their ends. As in solution, some laterally associated filaments were characterized by densities giving up the regular helical pitch 14 nm repeat giving rise to more open and unwound stretches. The co-incubation of GST-4xUb and LC3b-conjugated SUVs resulted in less dense network of filament bundles in comparison with GST-4xUb alone as well as SUV bound filaments (**Figure 3E**). Together, the cryo-ET visualization of purified p62 revealed the flexible properties of p62 filaments including cross-overs and branching, bundling of parallel p62 filaments in GST-4xUb bound condensates as well as the lateral association of the helical assembly on the surface of LC3b-conjugated membranes.

**Figure 3:**
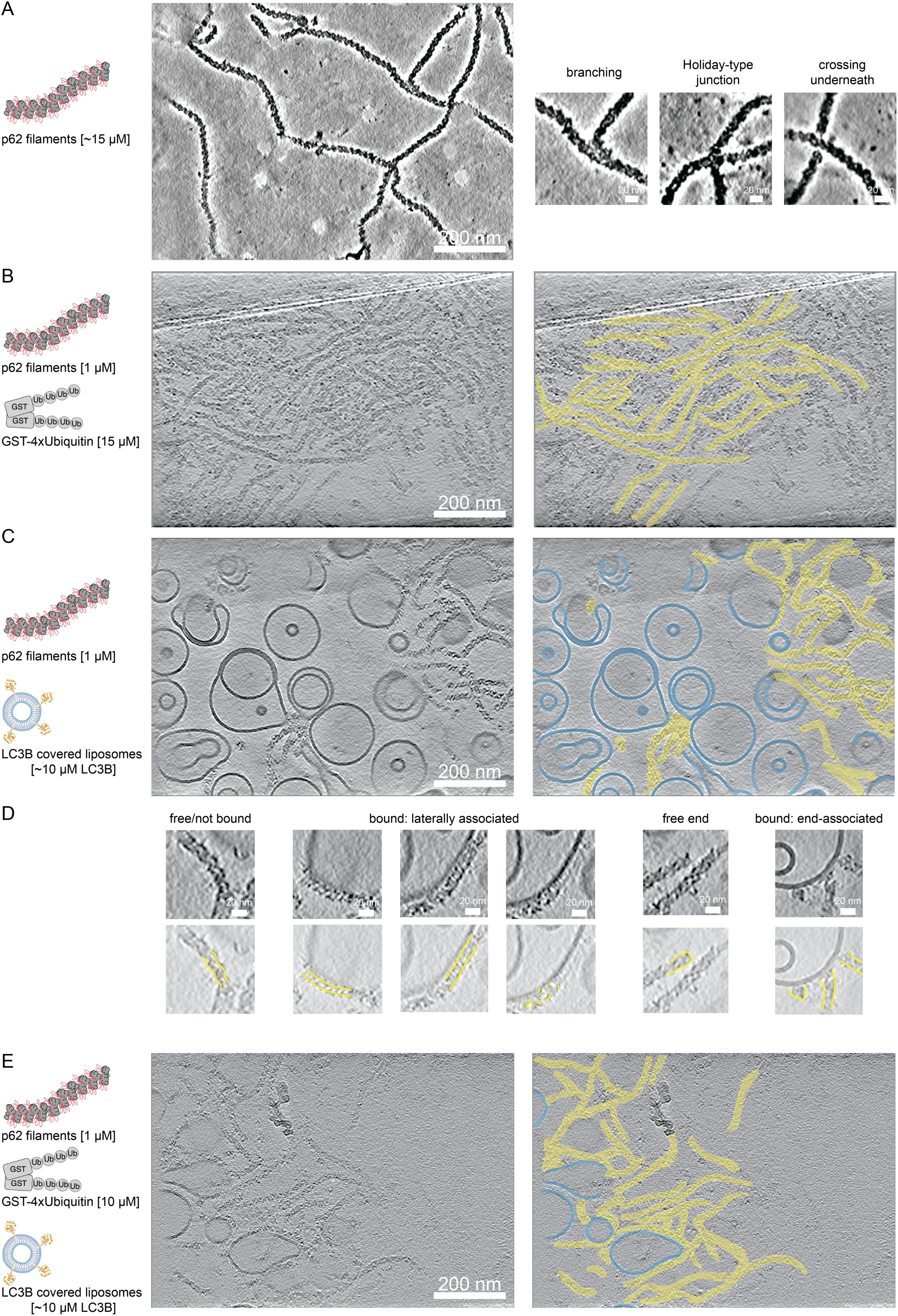
p62 filament interactions with LC3b-conjugated liposomes and poly-ubiquitin visualized by cryo-electron tomography. (A) Left. Representative micrograph of p62 filaments recorded with Volta phase plate. Gold fiducials were computationally removed in the locations of the smooth circles. Right. Close-up insets showing the regular 14 nm pitch features along the filament. p62 filament were found crossing each other underneath (left), branching by 90° (center) and cross overs as in the Holliday junctions of DNA (right). (B) Left. Upon addition of GST-4xUbiquitin, the formation of parallel p62 filament bundles was observed. Right. Segmentations of closely packed p62 filaments aligned in parallel. (C) The addition of pre-formed p62 filaments to LC3b-conjugated small unilamellar vesicles (SUVs) showed bound and unbound p62 filaments. (D) Top. p62 filaments were found not-bound (left), bound laterally associated with the membrane (left center), free ends (right center) or through their ends (right). Bottom. Outline of p62 densities is highlighted in yellow. For laterally associated p62, regular helical and unwound assemblies were observed. (E) Co-incubation of GST-4xUbiquitin and LC3b-conjugated SUVs led to a dense network of p62 bundled filaments partially bound to SUVs.

### ATG5 knockdown accumulates co-localized p62 punctae and lipid droplets

In order to further our understanding on the structural organization of autophagy receptor p62, we decided to visualize the cargo receptor in cells. Capturing rare and transient p62 punctae is challenging, therefore, we turned to a model system of ATG5 depletion that was shown to increase the size and number of p62 punctae in MEF cells and mice (Morishita et al., 2013; Pankiv et al., 2007) along with increased numbers of lipid droplets (Singh et al., 2009). We stalled autophagy by treating human RPE1 cells stably expressing mCherry-p62 with siRNA against ATG5 for 72 hours. To validate the phenotype upon ATG5 knockdown, we detected p62 punctae as well as lipid droplets by confocal fluorescence microscopy using mCherry-p62 and the lipid-droplet specific Lipi-Blue dye, respectively, in conditions of no transfection, mock transfection and transfection with ATG5 siRNA (**Figure 4A-C**). As a control for mCherry-p62, we imaged lipid droplets in RPE1 reference cells with Lipi-Blue after no transfection, mock transfection and transfection with ATG5 siRNA using confocal microscopy (**Figure 4D-F**). In order to confirm successful ATG5 siRNA knockdown, we performed a Western blot of the RPE1 cells showing a reduction in ATG5 expression levels by 90 % in comparison with the control samples (**Figure 4G**). When incubated with ATG5 siRNA, the number of p62 punctae was highest at an average of 37 per mCherry-p62 RPE1 cell (**Figure 4H**). In the presence of ATG5 siRNA, the average size of the lipid droplets increased from 0.30 to about 0.45 μm^2^ for mCherry-p62 RPE1 and for RPE1 cells (**Figure 4I**), while we also counted the highest number of co-localized p62 punctae in comparison with no or mock siRNA (**Figure 4J**). Interestingly, in our mock transfected cells we already observed a moderate increase in lipid droplet sizes and p62 punctae possibly due to the fact that cationic lipid mixtures have been observed to induce autophagy promoting LC3 conversion and p62 degradation (Man et al., 2010; Mo et al., 2012). Together with the autophagy-stalling effect of ATG5 siRNA, they likely contribute to further enrichment of p62 positive structures. In conclusion, when transfecting RPE1 cells with ATG5 siRNA, we observed an accumulation for p62 punctae and lipid droplets as well as a size increase for lipid droplets. The characterized condition of ATG5 knockdown may be well suited for the in situ visualization of p62 punctae as p62 cargo uptake in the lysosome is impaired.

**Figure 4:**
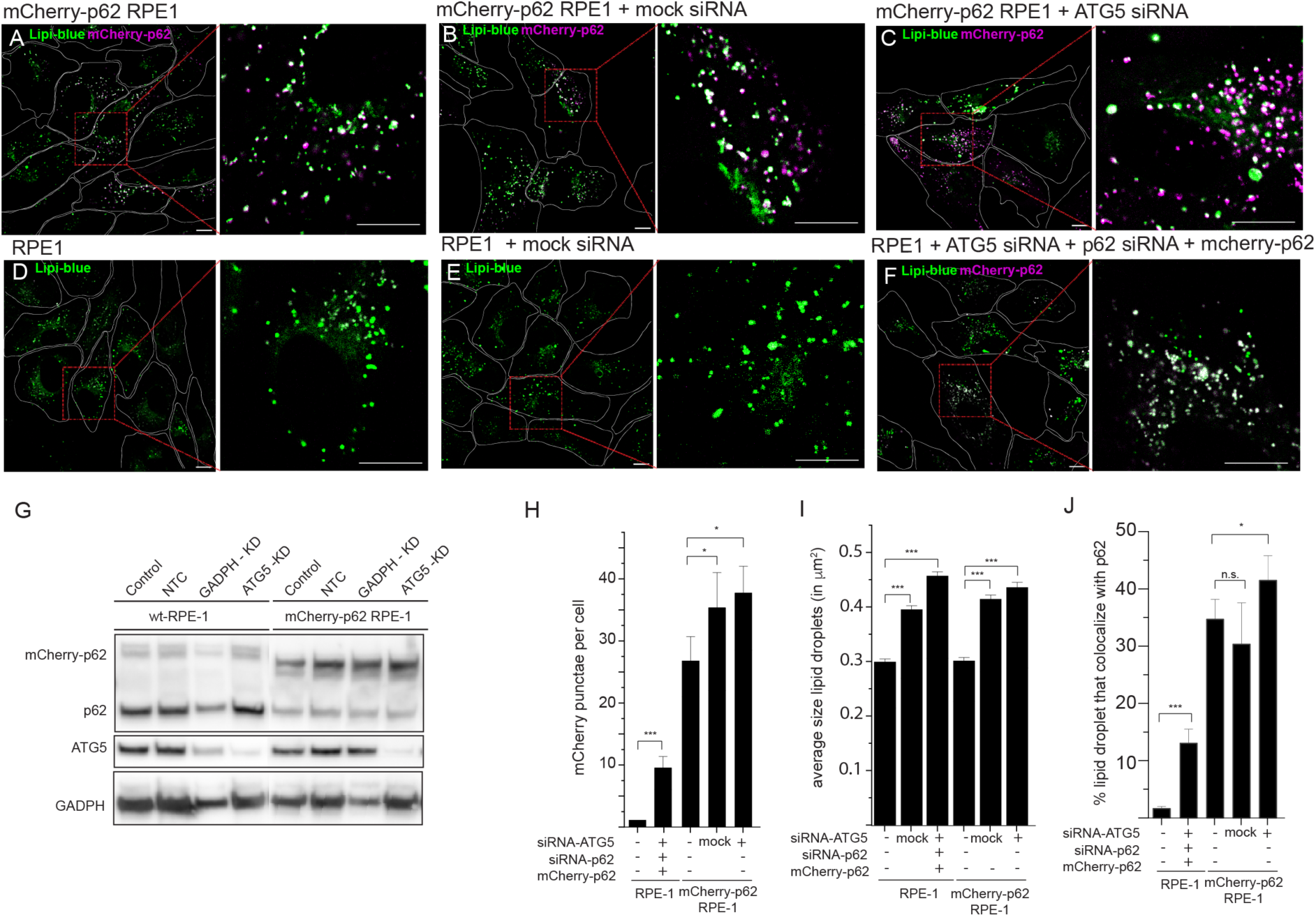
Confocal fluorescence microscopy of p62 punctae and lipid droplets upon ATG5 knockdown. mCherry-p62 RPE1 cells stained with Lipi-Blue for 24 hours at conditions of (A) no transfection, (B) 72 hrs after a mock transfection with RNAiMax and (C) 72 hrs after a transfection with ATG5 siRNA. Reference RPE1 cells stained with Lipi-Blue for 24 hours at conditions of (D) no transfection, (E) 72 hrs after a mock transfection with RNAiMax and (F) 72 hrs after a transfection with ATG5 siRNA. (G) Western blot confirming the knockdown of ATG5 expression including a control knockdown of GADPH. Quantifications for the displayed samples: (H) Number of p62 structures per cell, (I) size of lipid droplets and (J) number of p62 punctae and lipid droplets colocalizations. Statistical significance was determined with a t-test: *=p<0.1, =p<0.01,* =p<0.001

### p62 encapsulates lipid droplets by a discontinuous coat of variable thickness

Based on the established condition of accumulated colocalized p62 punctae and lipid droplets, we set out to perform detailed in situ structural analyses to study p62 during lipid droplet cargo recognition. The RPE1 cells were grown under the same conditions described above and placed on micropatterned electron microscopy grids, vitrified, subsequently thinned to ∼150 nm using a focused ion beam (FIB) and visualized using a scanning electron microscope (SEM) (**Figure 5A**). To verify that the lamella contained mCherry-p62, we imaged the sample using an integrated cryo-fluorescence microscope and correlated it with the FIB-SEM image (Smeets et al., 2021) (**Figure 5B**). When we visualized these p62 structures using super-resolution cryo-confocal microscopy, they showed circularly resolved mCherry-p62 signal on their rim suggesting the cargo encapsulation by the autophagy receptor p62 (**Figure 5C**). At these sites, a total of 25 tomograms were recorded while those with green autofluorescence from lysosomes were not further analyzed. In the reconstructed tomograms, we identified characteristic structures made of a homogeneous texture of spherical shape suggestive of a lipid droplet (Ganeva et al., 2023) (**Figure 5D**) in agreement with the frequent co-localization of p62 and lipid droplet punctae demonstrated above. A dark and highly dense layer was found surrounding the droplet in addition to a single phospholipid bilayer. The homogeneously textured spheres were similar in size measuring between 510 - 530 nm (**Figure 5E**). The surrounding layer, or coat, varied in thickness within a single structure as well as between structures ranging between 4 and 27 nm, with a mean of 12.6 nm (**Figure 5F** and **Supplementary Figure 4**). The distance between the surface of the sphere and the surrounding single lipid membrane was relatively uniform at 13.8 nm with some weaker protein densities filling this space while other densities connecting these two structures (**Figure 5G/H**). The segmented tomograms revealed that when the layer was thin, patches or islands of dark contrast were found distributed around the surface of the lipid droplet (**Supplementary Figure 5**). However, when the layer was thick, it was completely continuous. Next to the lipid droplet, several small vesicles between 35 and 52 nm in diameter were observed consistent with previously described ATG9A vesicles (Mailler et al., 2021). In summary, ATG5 knockdown in human RPE1 cells yielded p62 positive spherical structures of 500 nm diameter composed of an inner lipid droplet and a p62-containing, dense layer of variable thicknesses covering the surface surrounded by a single bilayer membrane.

**Figure 5:**
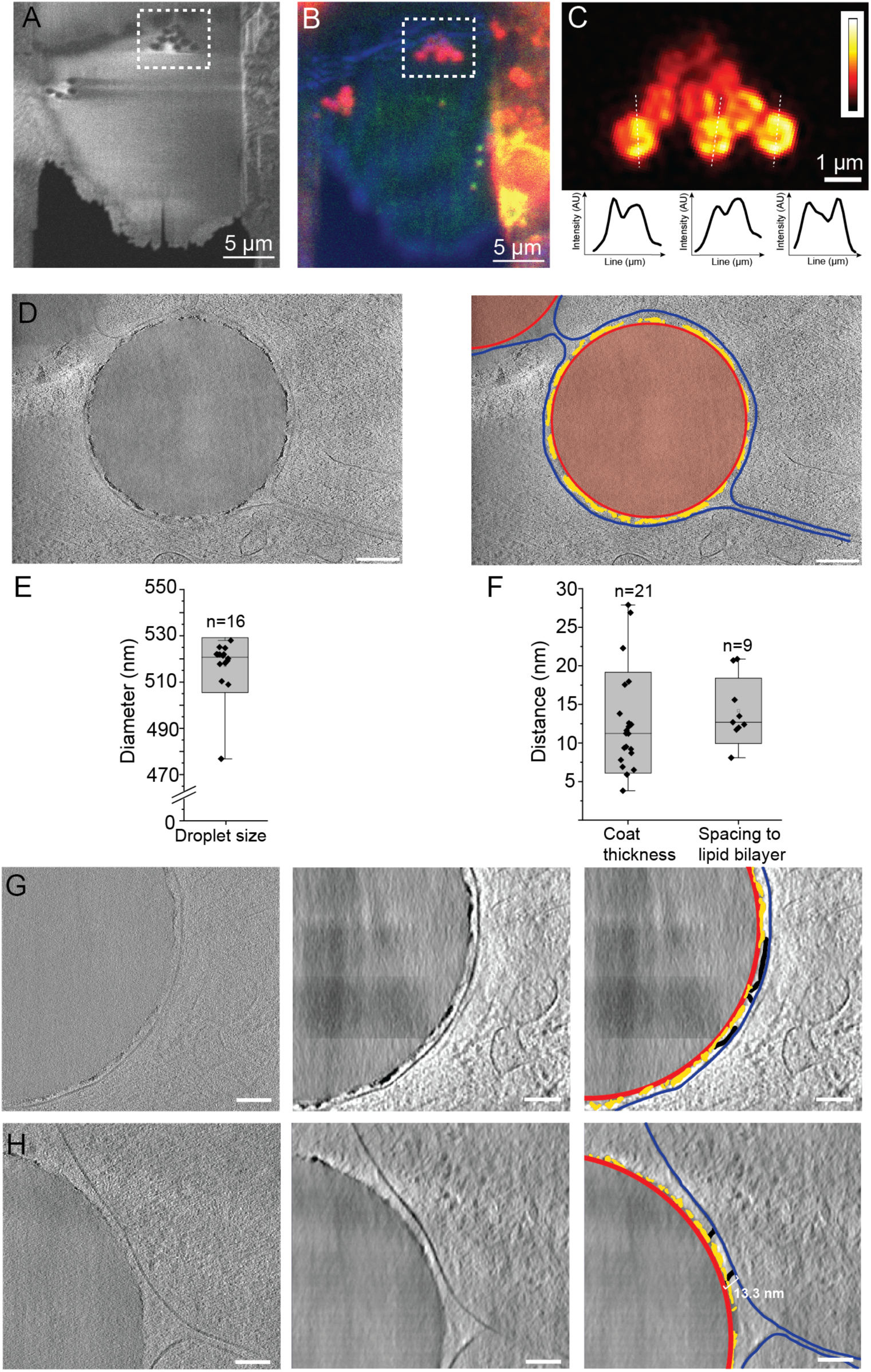
Ultrastructural in situ characterization of p62-encapsulated lipid droplets. (A) Scanning electron microscopy image of a lamella generated from an EM grid grown with RPE1 cells. (B) Correlated fluorescence image of the corresponding lamella recorded with a reflected light channel and fluorescent channels of 595 nm for mCherry-p62 and 515 nm for autofluorescence. (C) Super-resolution cryo-confocal fluorescence image recorded at 561 nm for mCherry-p62 showed three p62-positive structures resolving a circular fluorescence signal as displayed in three corresponding intensity profiles below. (D) Tomographic gray-scale z-slice through the homogeneous density sphere surrounded by a dark high-contrast layer with an enwrapped single membrane. Scale bar 100 nm. (E) Segmented and labeled structure superimposed on gray-scale z-slice, light red: lipid droplet with red: droplet surface, yellow: dark and high-contrast surrounding p62 layer and blue: single lipid membrane. (F) Left. Box plot indicates a well-defined lipid droplet diameter of about 520 nm. Right. Box plot of measured distances of observed dark layer thickness and spacing of droplet surface to enwrapped single lipid bilayer. (G) Tomographic z-slice of half a lipid droplet in raw grayscale (left), denoised grayscale (center) and superimposed labeled structures of droplet surface (red), dark layer (yellow), weaker protein density (black) and enwrapping single lipid bilayer (blue). Scale bar 50 nm. (H) Tomographic z-slice of another droplet showing the spacing of 13.3 nm between droplet surface and single lipid bilayer. Scale bar 50 nm.

### Elemental analysis reveals calcium and phosphorus in the p62 positive layer

Given the strong observed intensity of the surrounding p62-positive layer with respect to the grey levels of the lipid droplet and cytosol, we wondered whether non-organic elements contributed to the high contrast. To address this question, we employed scanning transmission electron microscopy (STEM) coupled with energy-dispersive X-ray (EDX) spectroscopy to record element-specific X-rays from the mCherry-p62 RPE1 lamella treated with ATG5 siRNA investigated above (**Figure 6A-D**). For this sample, we acquired STEM images on a high-angle annular dark-field (HAADF) detector and simultaneously recorded the back-scattered EDX signal (**Figure 6E**). The EDX signal was averaged over the excised regions of the cytosol, lipid droplet core and the p62 coat (**Supplementary Figure 6**). For the cytosol, we detected mostly carbon (C), nitrogen (N) with low and oxygen (O) with high intensity at approx. 300, 400 and 500 eV, respectively. For the lipid droplet, we observed enriched C in comparison with the cytosol, reduced N and O consistent with the presence of triacyl glycerides and sterols within the sphere of lipid droplets. In both regions, no inorganic elements were observed at higher energies. For the p62 coat, we found the C, N, O signal comparable to the cytosol with clear additional inorganic signals of magnesium (Mg), calcium (Ca) and phosphorus (P) at 1250, 2000 and 3800 eV, respectively, supporting the observation of the relatively strong contrast in the raw tomograms. The detected high levels of Ca and P suggest the presence of calcium phosphate. As a control, we focussed on lipid droplets showing no p62 colocalization and obtained a very similar EDX signal (**Supplementary Figure 7**). Other p62 positive single-membrane vesicular structures resembling lysosomes showed no differences in EDX profiles between the vesicular area and the cytosol (**Supplementary Figure 8**).

**Figure 6:**
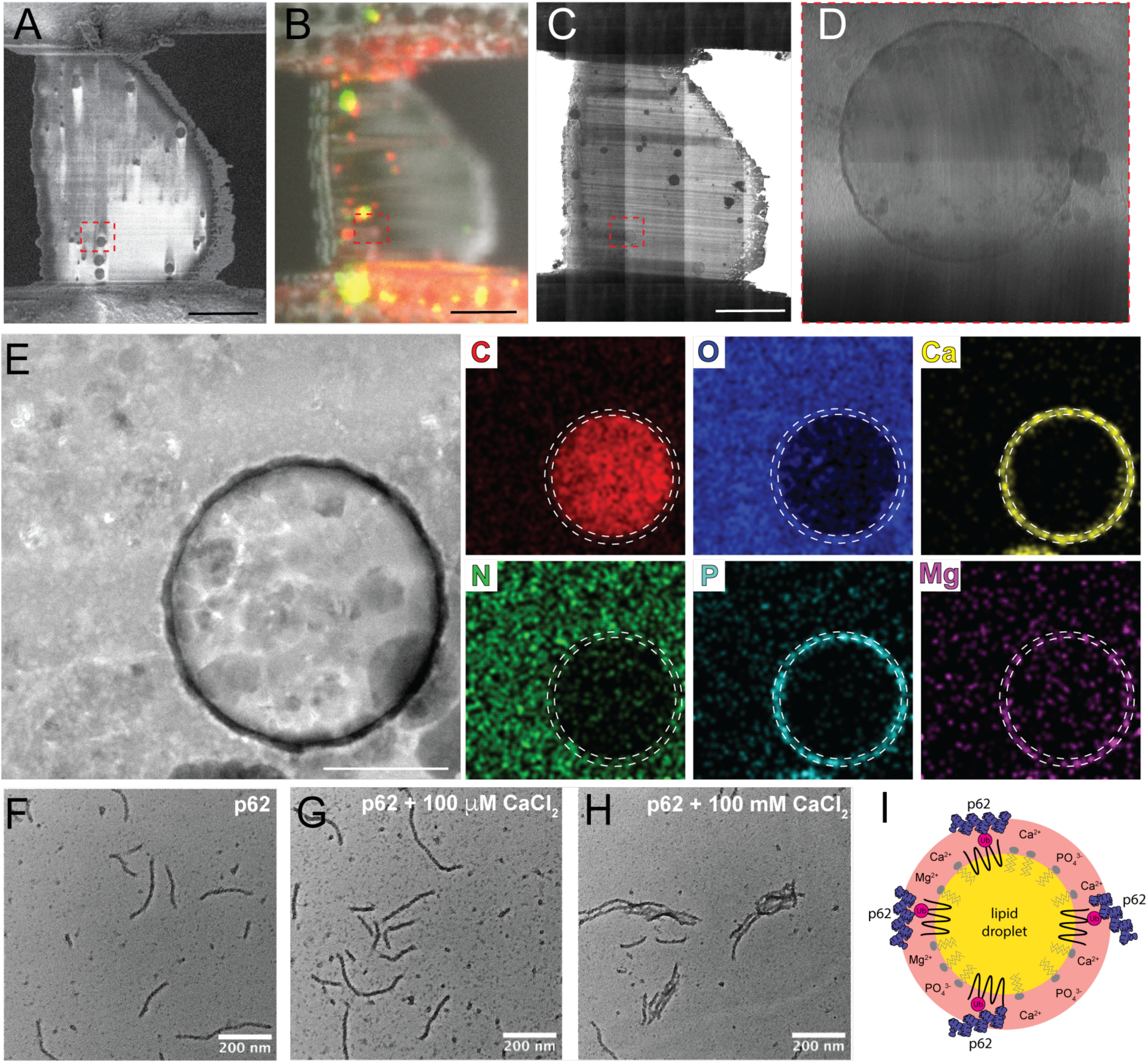
Compositional elemental analysis of p62-encapsulated lipid droplets using energy dispersive X-ray (EDX) spectroscopy. (A) Scanning electron microscopy (SEM) image of milled lamella with inset of characterized lipid droplet. (B) Correlated SEM image superimposed with a reflected light channel and fluorescent channels of 595 nm for mCherry-p62 and 515 nm for autofluorescence. (C) Transmission electron microscopy image of corresponding lamella. (D) Enlarged image of p62-positive lipid droplet. (E) High-angle annular dark-field scanning transmission electron microscopy (HAADF-STEM) image of droplet of interest with corresponding EDX signals for Carbon in red (C), Oxygen in blue (O), Calcium in yellow (Ca), Nitrogen in green (N), Phosphorus in cyan (P) and Magnesium in purple (Mg). Scale bar 500 nm. Negative stain EM images of (F) purified p62 filaments, (G) with 100 µM CaCl_2_ and (H) 100 mM CaCl_2_. (I) Model summary of analyzed p62 droplet structures: the lipid droplet devoid of any phospholipids is encapsulated by p62-positive layer that is rich in CaPO_4_.

Given the unexpected finding of inorganic elements located in the p62 coat, we tested whether the addition of Ca^2+^ ions had an effect on purified p62 filaments. When increasing the CaCl_2_ concentrations from 100 nM to 100 mM, we observed a bundling of p62 filaments at 100 mM CaCl_2_ in negative staining EM (**Figure 6F-H**). Moreover, when testing other divalent cations such as Zn^2+^ and Mg^2+^, we found that the addition of ZnCl_2_ had a stronger clustering effect already visible at 100 µM, while the addition of MgCl_2_ did not show a significant bundling (**Supplementary Figure 8**). Together, these experiments show that the observed p62-positive structures correspond to lipid droplets surrounded by a coat made of p62, including significant amounts of calcium (**Figure 6I**) that also promotes the formation of p62 oligomer bundles in vitro and may explain the characteristic dense layer observed in the tomograms.

## Discussion

In the current study, we addressed the structural organization of the archetypical autophagy receptor p62/SQSMT1 at various levels of resolution. Initially, we determined the cryo-EM structure of full-length p62 at 4.5 Å global resolution revealing the double helical filament PB1 scaffold with peripheral lower-resolution densities corresponding to the ZZ-domain and C-terminal part (**Figure 1**). Interestingly, the addition of LC3b led to a shortening of the filament at higher concentrations while GST-4xUb resulted in the bundling of filaments to phase-separated entities consistent with solvent-exposed accessible binding sites to LC3b and ubiquitin through the major groove of the filament (**Figure 2**). When p62 filaments were bound to LC3b-conjugated SUVs, we observed a lateral association on model membranes of the p62 filament while maintaining regular p62 oligomer arrays (**Figure 3**). Moreover, we visualized p62 structures in cells using fluorescence light microscopy, correlative light and electron microscopy (CLEM) followed by cryo-ET in a p62-enriched phenotype due to ATG5 knockdown (**Figure 4 and 5**). The detailed ultrastructural analysis of this autophagy-stalled phenotype revealed a high-contrast layer of p62-positive structures surrounding lipid droplets due its enriched calcium phosphate content (**Figure 6**).

In contrast to the previously determined PB1 assemblies (Jakobi et al., 2020), the here determined full-length p62 filament structure formed a double-helical filament architecture widely known from the classical DNA double helix model (Watson & Crick, 1953). The full-length structure revealed two well-resolved antiparallel PB1 scaffold strands while the PB1 domain-only assemblies have one or two additional PB1 strands inserted into the helical assembly leading to an almost closed tube. In the full-length structure, the space required by the additional 340 C-terminal amino acids restricts the assembly to two antiparallel PB1 strands leaving space for a major groove. Our binding assays showed a shortening the filament upon LC3b interaction in agreement with model predictions of the LIR motif to be accessible in solution through the major groove. In our solution studies, we also found the p62 concentrations to be critical for detecting clear binding effects such as shortening or filament bundling in the case of 4xGST-Ub. In addition, we also observed bundling of p62 filaments upon addition of divalent cations in CaCl_2_ and ZnCl_2_ solutions (see **Figure 6G-H**). Multivalency available in a polymer and across polymers may mediate the bundling, e.g., when several UBA domains from different filaments bind to polyubiquitin or when parts of ZZ-domains originating from multiple p62 filaments share the coordination of divalent cations such as Zn^2+^ and Ca^2+^. Multivalent interactions mediated by GST4xUb have been shown to promote phase separation of p62 (Zaffagnini et al., 2018a). In the cell, we expect some of these interactions to be present on membrane surfaces through membrane-linked p62 polymers and oligomers, which provides the principal benefit of high local concentrations offered for low-affinity interactions to be enhanced by high avidity binding. In support of this concept, when we linked the binding partner LC3b (Pankiv et al., 2007) to the liposome membrane surface, upon p62 filament addition we did not observe a complete filament disassembly, rather we observed a lateral binding of the filament. In the images at physiological pH, we also found an unwinding and structural opening up of the double-helical filament architecture located on the membrane surface. In such an assembly configuration, previously inaccessible available binding sites in the interior will become accessible in an opened p62 “bubble” structure while still maintaining functional polymers and oligomers. The unwinding, structural opening and re-arrangement of the filament architecture on membrane surfaces may provide a higher level of regulating access of binding partners to the multiple binding sites in p62.

After the in vitro structural and biochemical work, we turned to the RPE1 cellular model to investigate the ultrastructural details of p62 using cryo-ET. In order to enrich p62 structures, we depleted ATG5 that is an essential member of the core machinery driving autophagy progression (Mizushima et al., 2001; Morishita et al., 2013; Pankiv et al., 2007). During autophagy, ATG5 is covalently linked to ATG12 that together with ATG16L is required for LC3-I conjugation to the membrane in the form of LC3-II and thereby accomplishing phagophore expansion (Otomo et al., 2013). Upon ATG5 depletion, characteristic LC3 punctae and autolysosomes cannot form leading to an accumulation of cargo, e.g., when lipid droplet breakdown is inhibited, triacylglycerides have been shown to accumulate (Singh et al., 2009). Moreover, in mice, ATG5 knockouts have severe consequences for neuronal development leading to delays in maturation as ATG5 appears to be essential for the survival of adult-generated neurons (Xi et al., 2016). Based on the near-native visualization using in situ cryo-CLEM in RPE1 cells, we observed p62-positive structures wrapping around lipid droplets made of triacylglycerides giving rise to homogeneous density in the electron tomograms and characteristic EDX signals. Lipid droplets have been observed in the diameter ranges of 100 nm to over 1 μm in size (Schott et al., 2019) while the here observed ones had relatively narrow distribution of 500 to 540 nm. Lipophagy has been described as being initiated by the ubiquitination of PLIN proteins embedded in the phospholipid monolayer surrounding the lipid droplets (Tsai et al., 2017). Moreover, LC3 and p62 were also described as forming punctae on the surface of lipid droplets (Robichaud et al., 2021). In this way, they are thought to induce the formation of the autophagosomal membrane surrounding the lipid droplets. While in ATG5 depleted cells LC3-II is not present on the membrane, p62 has been shown to still be capable of recruiting cargo, ATG9A vesicles, ATG16L vesicles and FIP200 condensates (X. Feng et al., 2023; Itakura & Mizushima, 2011). Possibly, the small vesicular structures we observed next to the p62 positive lipid droplets (see **Supplementary Figure 5B**) may correspond to previously observed ATG9 vesicles (30-60 nm) or ATG16L vesicles (Dupont et al., 2014; Mailler et al., 2021; Puri et al., 2013; Yamamoto et al., 2012).

When we segmented tomography data of the ATG5 depleted cells, we observed p62 coating of various thicknesses on the lipid droplets. In close vicinity, we often found a single, unclosed membrane (see **Supplementary Figure 5A**) enwrapping the coated lipid droplet. In the ATG5 knockdown condition, we do not expect a bona fide autophagy double membrane to be present. Due to the observed connections to membrane stacks, it appears plausible that we visualized recruited ER membrane unable to extend and seal off the lipid droplets due to the lack of lipidated LC3-II. In our p62-positive structures, we also observed 13.3 nm wide protein density connecting the droplet surface/coat and the enwrapped membrane in addition to elongated flexible density. The density is not compatible with regular helical p62 filaments used for structure determination above as no pitch repeat distance along the long axis is discernible. Alternative assemblies, however, such as unwound or opened p62 filaments share similar width dimensions to the measured 13.3 nm. Other potential molecules of the autophagy core machinery include ATG2A/B (Y. Wang et al., 2024), the ULK1 complex (Chen et al., 2023) or FIP200 (Chen et al., 2023) all of which have elongated shapes but are either longer or shorter than the observed densities. Clearly, more research and improved imaging features will be required to confirm the identity of the protein complexes involved in the studied p62-positives structures.

The p62-lipid droplet structures we have visualized in this study were found surrounded by dark layers. We identified them to be mainly composed of calcium, phosphorus and magnesium using EDX spectroscopy suggesting high levels of calcium phosphate. Previously, the same approach has been used on vitreous sections and other frozen hydrated samples (Kumar et al., 2020; WOLF et al., 2015; Wolf et al., 2017; Zierold et al., 1984) while in this study we employed it on FIB-milled lamella. A dark high-contrast layer had been observed before in close proximity to lipid droplets in the context of autophagy using plastic section electron microscopy (Robichaud et al., 2021; Schott et al., 2019; Singh et al., 2009). We do not claim that the previously imaged structures correspond to structures we observed, nevertheless the resemblance is noteworthy. For some time, intracellular calcium levels have been linked to autophagy as rapamycin-induced autophagy led to an increase in cytosolic Ca^2+^ concentration and a decrease in the calcium ER leak rate both of which were independent of LC3 lipidation (Decuypere et al., 2013). Moreover, the addition of a calcium chelator such as BAPTA-AM was shown to be sufficient to block autophagy initiation (MacVicar et al., 2015). In a more recent study, evidence was provided that brief calcium transients are released from the ER, which trigger FIP200 punctae formation and modulate phase separation at the autophagosome assembly site (Q. Zheng et al., 2022). In yeast, calcium has also been shown to be required for autophagy initiation under glucose starvation conditions (Yao et al., 2024). It seems plausible that in this study we directly observed calcium ions bound to the lipid droplet surface originally released from the ER, which help to organize the early stages of lipophagy. Given the demonstrated bundling capabilities of p62 by divalent cations in solution, it is possible that the observed dark p62-positive coat found on the cargo lipid droplet is the result of calcium transients release from the ER that orchestrate the early stages of autophagy machinery recruitment. Further research will be required to fully establish the links between the here presented cryo-EM structure, biochemical binding properties, higher-order organization, in situ ultrastructure and autophagy initiation.

## Supplementary Figures

**Supplementary Figure 1:**
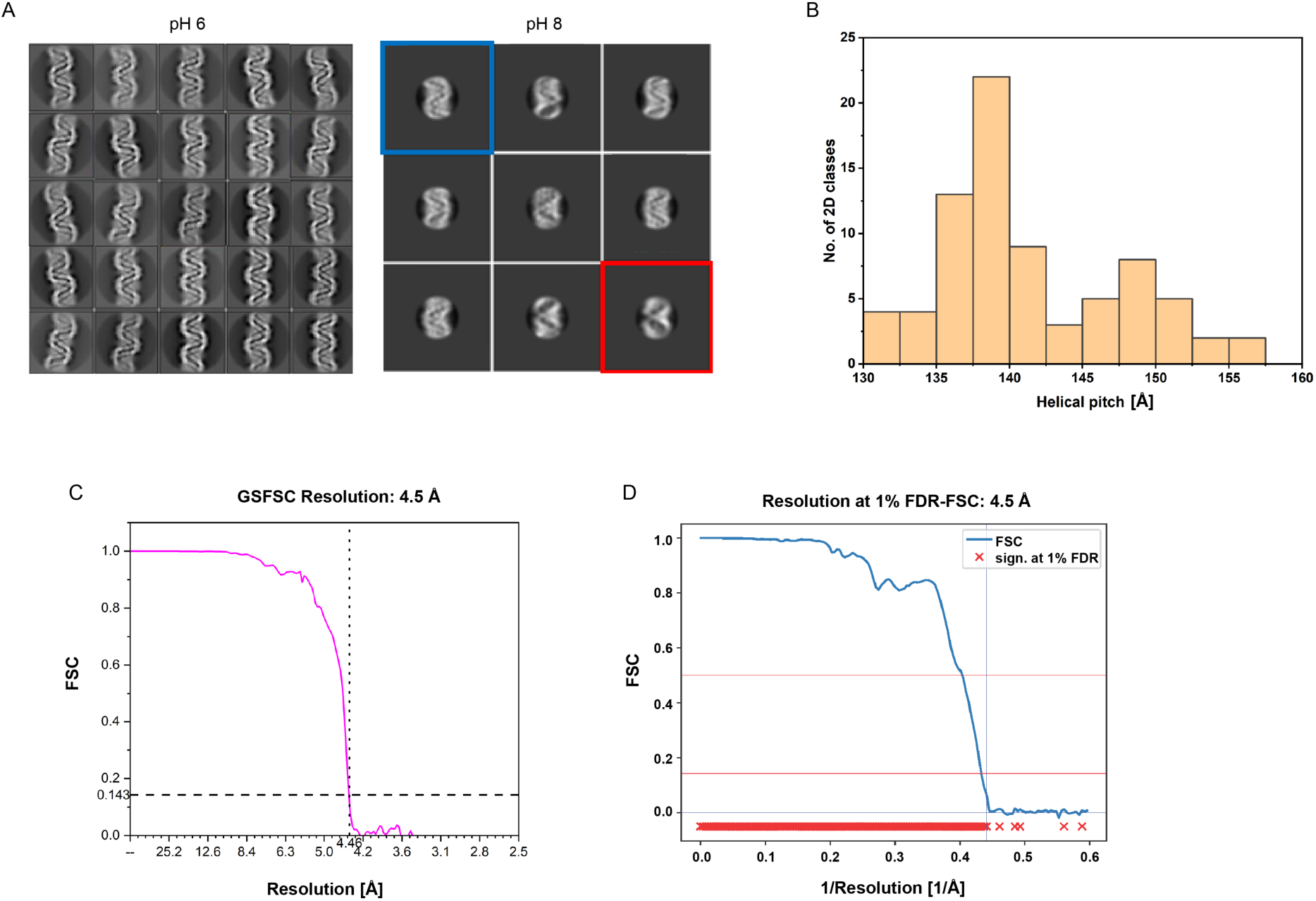
p62 filament sample optimization, helical pitch of p62 filaments and resolution assessment of p62 cryo-EM structure. (A) 2D classification at pH 6 (left) and pH 8 (right). At pH 6, the classes show a regular helical architecture. At pH 8 almost 60% of the particles show an unwinding of the two strands (red), whereas 40% of the particles still show a closed form of the filament (blue). (B) Helical pitch distribution of p62 filaments ranges from 130 to 160 Å. (C) Fourier shell correlation (FSC) curve of the p62 filament density map indicates a resolution according to the cutoff FSC(0.143)=4.5 Å. (D) FDR-FSC curve indicates a resolution of 4.5 Å.

**Supplementary Figure 2:**
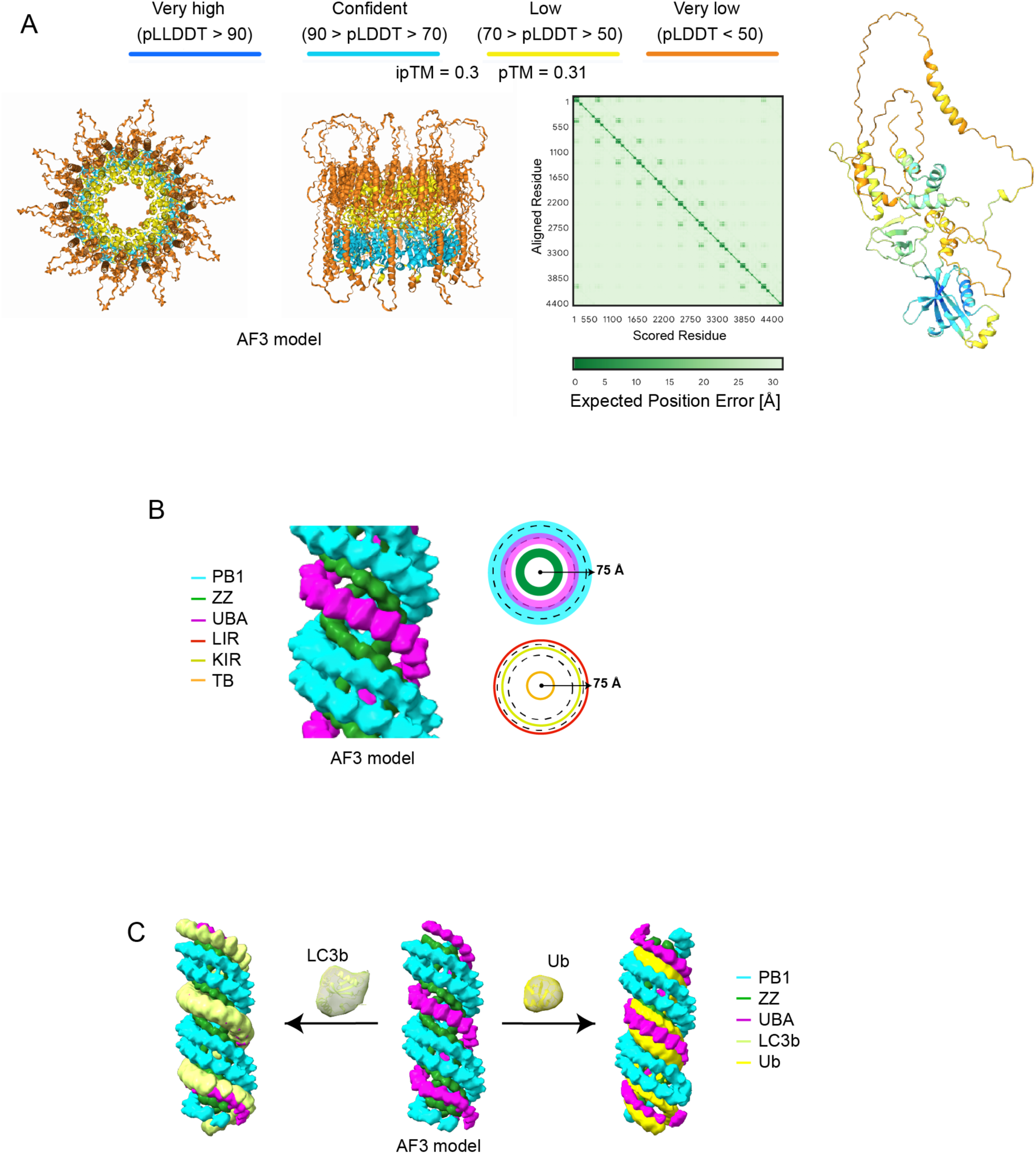
AlphaFold3 prediction of p62 assembly, domain localization, binding of LC3b and competition with ubiquitin. (A) Left. AlphaFold3 (AF3) model of the p62 decamer ring (top and side view), colored according to a per-atom confidence estimate. Predicted aligned error (PAE) plot reveals spatial proximity of the domains. Right. Single p62 molecule AF3 prediction excised from 10-mer colored with per-residue model confidence estimate. (B) p62 filament model of AF3 predictions showing PB1 (cyan), ZZ (green) and UBA (magenta) domains rendered at 15 Å resolution. Top views showing relative position of subunits and binding sites and their radial distance from the center of the filament. The two dotted lines mark the position of the PB1 domain that corresponds to the strongest observed density in the cryo-EM map. (C) p62 filament models of AF3 predictions rendered at 15 Å resolution with LC3b (left) or ubiquitin (right) bound.

**Supplementary Figure 3:**
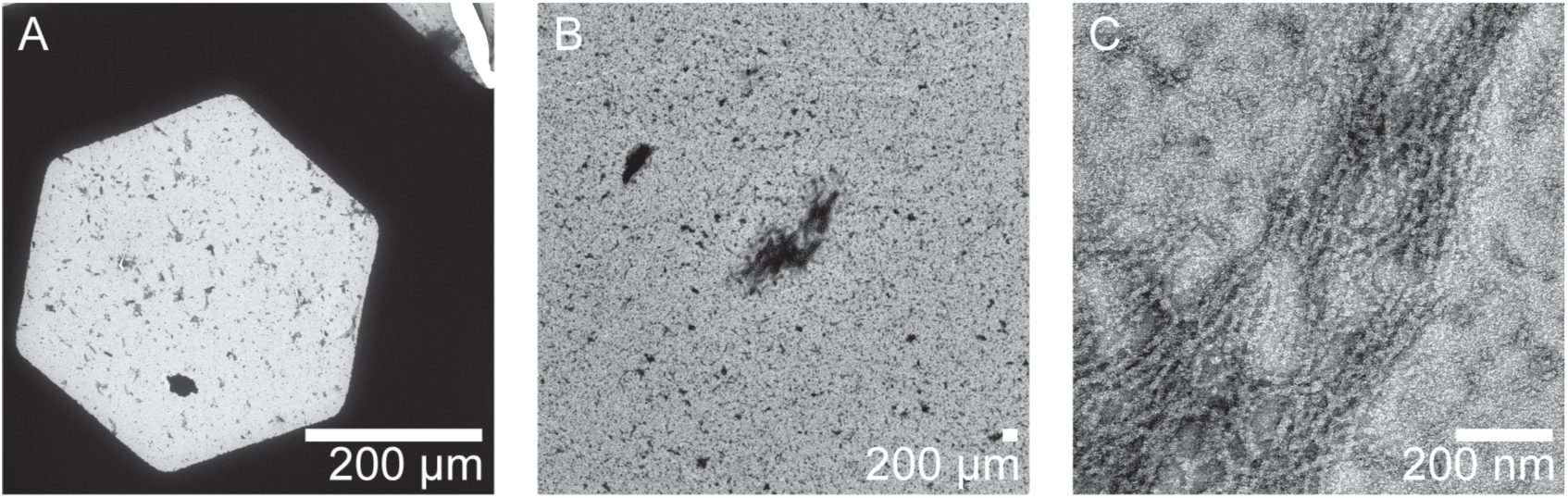
Phase separation of p62 filaments induced by addition of GST-4xUbiquitin. p62 filaments at 1 µM concentration were mixed with 15 µM GST-4xUbiquitin, incubated for four minutes and subsequently negatively stained. The resulting phase separation droplets were imaged at (A) 100x, (B) 8,500x and (C) 57,000x magnification revealing dense clusters of p62 filament bundles.

**Supplementary Figure 4:**
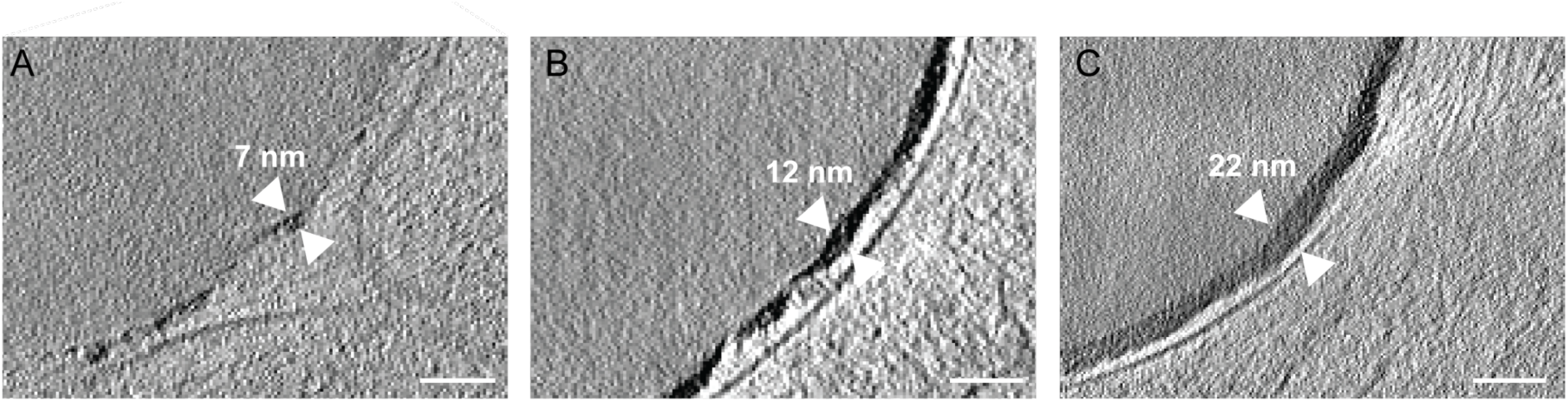
Different thicknesses of droplet coats. Lipid droplets are coated by a high-contrast layer of different thicknesses measuring (A) 7 nm, (B) 12 nm and (C) 22 nm. Scale bar 50 nm.

**Supplementary Figure 5:**
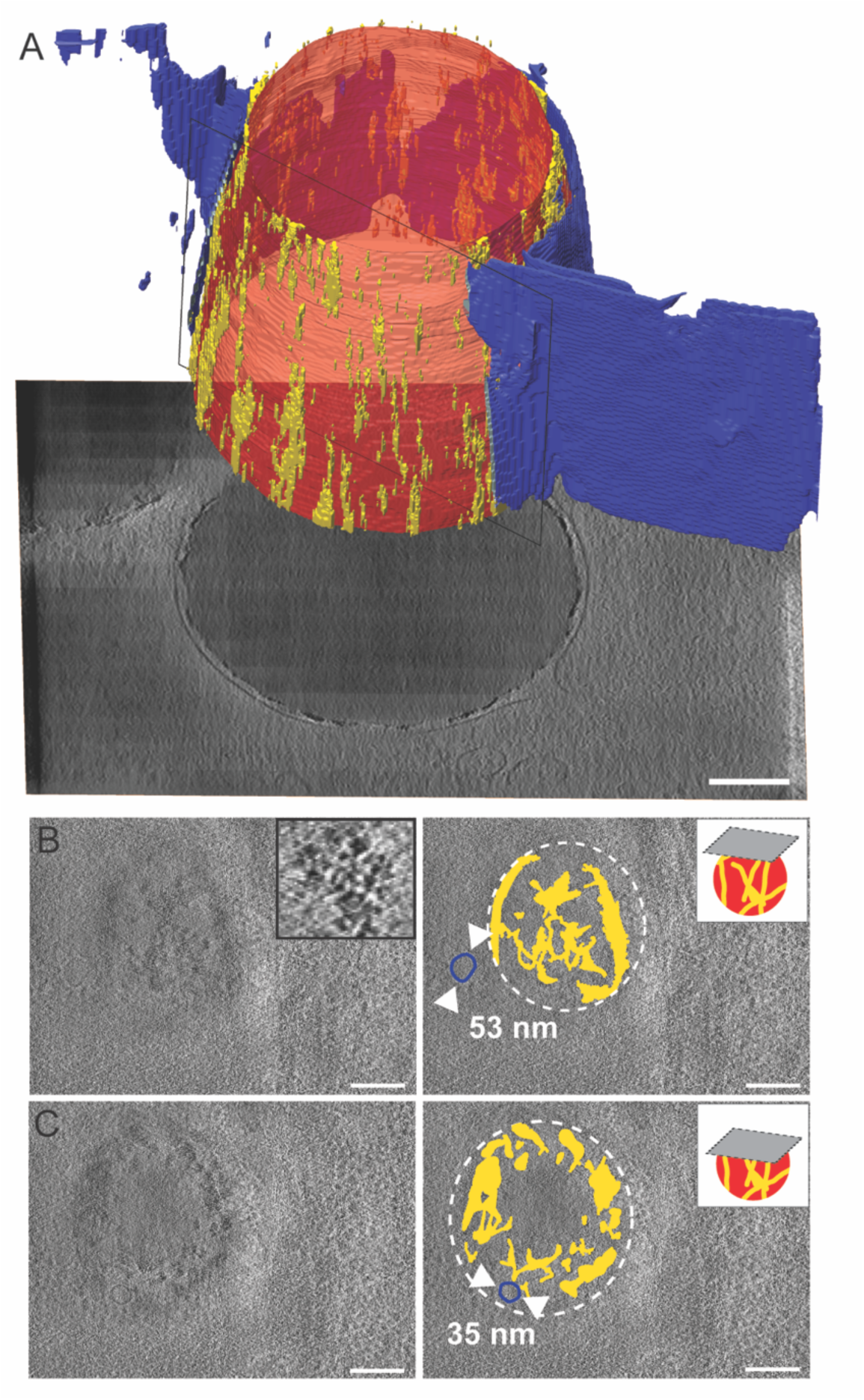
Segmented tomogram showing a thin discontinuous high-contrast p62-positive coat surrounding the lipid droplet. (A) Segmented tomogram rendered in three dimensions: lipid droplet (red), p62-positive coat (yellow) enwrapped by a single lipid bilayer stack (blue). The membrane was opened up using a clipping plane (black rectangle) for easier visualization. (B) Upper z-section through segmented tomogram reveals the thin discontinuous surface coating around the lipid droplet. Inset shows the granular nature of the surface coating. (C) Center z-section of the same lipid droplet. Small vesicular structures (blue) of 35 and 53 nm diameter are also visible in both z-slices. Scale bar 100 nm.

**Supplementary Figure 6:**
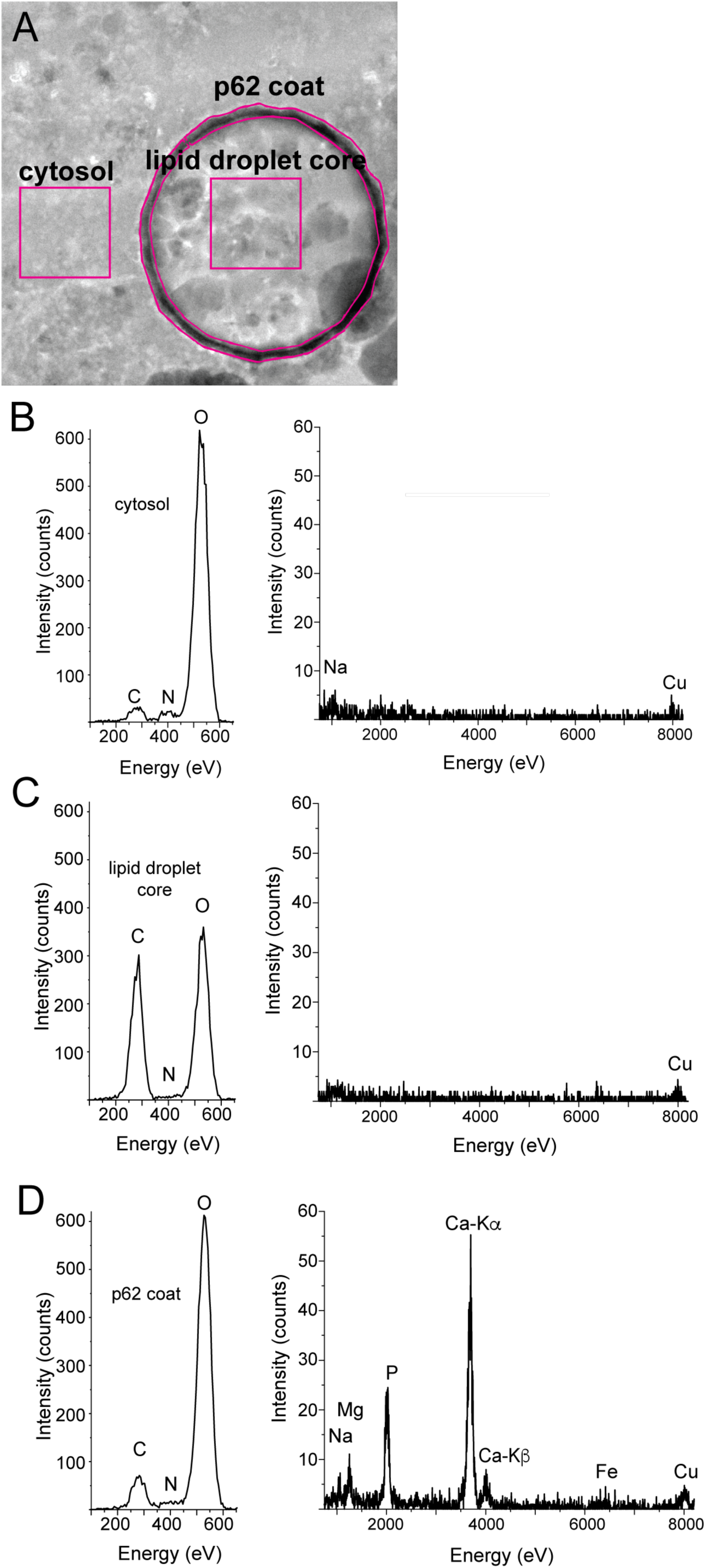
Energy-dispersive X-ray (EDX) spectroscopy combined with STEM imaging of p62-positive lipid droplets. (A) HAADF-STEM image of p62 positive lipid droplets. Insets indicate the regions used for summing of EDX signals. The same number of pixels were summed in all regions. X-ray spectra showing which elements are present in the different areas; in (B) cytosol, (C) lipid droplet core and (D) dense p62 positive coat, respectively. Peaks were marked with their respective elements. The weak copper (Cu) peak originates from the copper sample holder used.

**Supplementary Figure 7:**
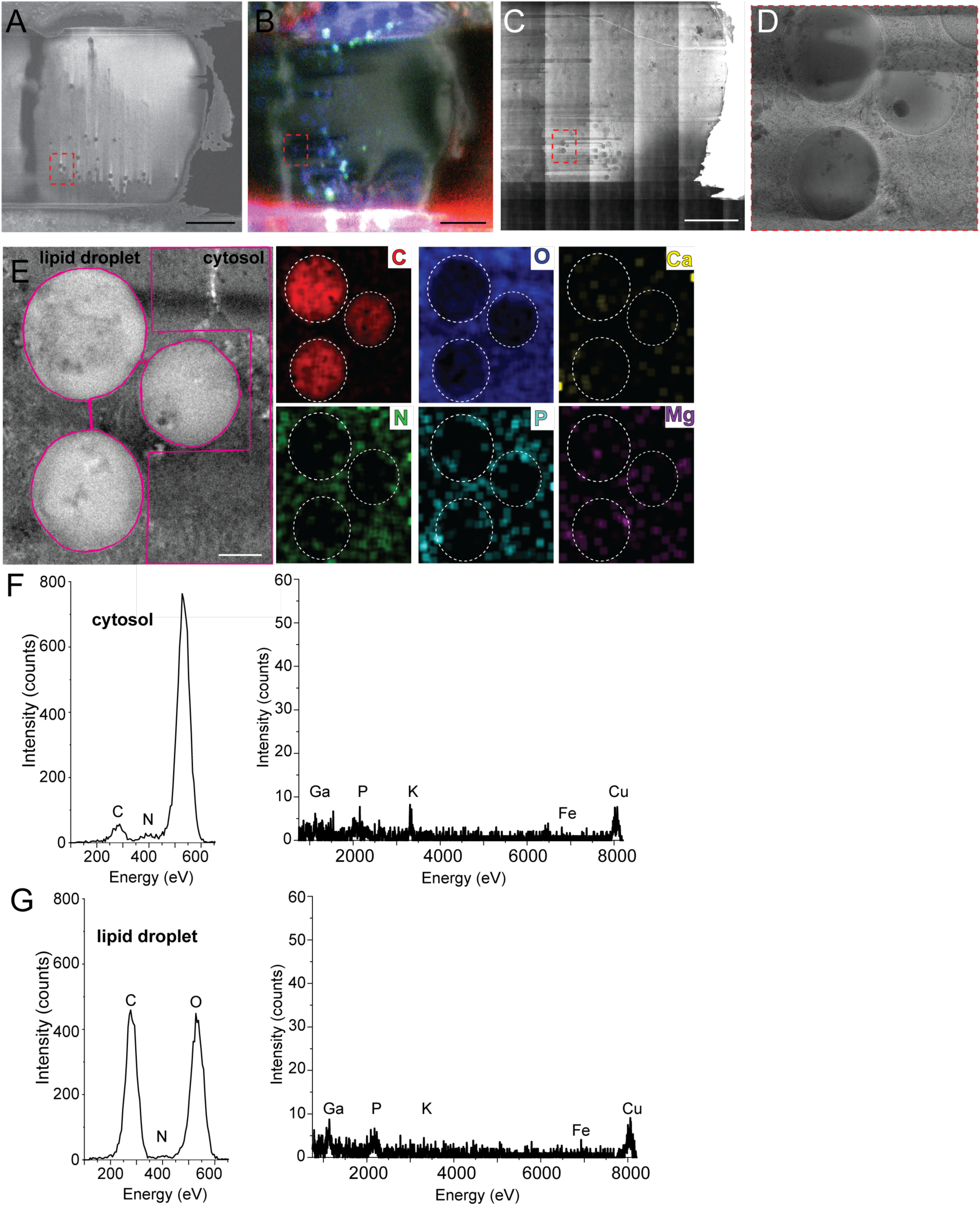
Energy-dispersive X-ray spectroscopy (EDX) combined with STEM imaging of native lipid droplets. (A) Scanning electron microscopy images of lamella with visible droplets in inset. Scale bar 5 µm. (B) SEM image superimposed with fluorescent image of Lipi-Blue dye. (C) Overview image of corresponding recorded transmission electron microscope (TEM). (D) Three spatially close lipid droplets in higher magnification TEM image. (E) HAADF-STEM image of three adjacent lipid droplets corresponding elemental images of carbon (C), oxygen (O), calcium (Ca), nitrogen (N), phosphorus (P) and magnesium (Mg). The image shows two insets of the cytosol and droplet region used for subsequent averaging. EDX spectra of (F) cytosol region and (G) lipid droplet regions. Scale bar 200 nm.

**Supplementary Figure 8:**
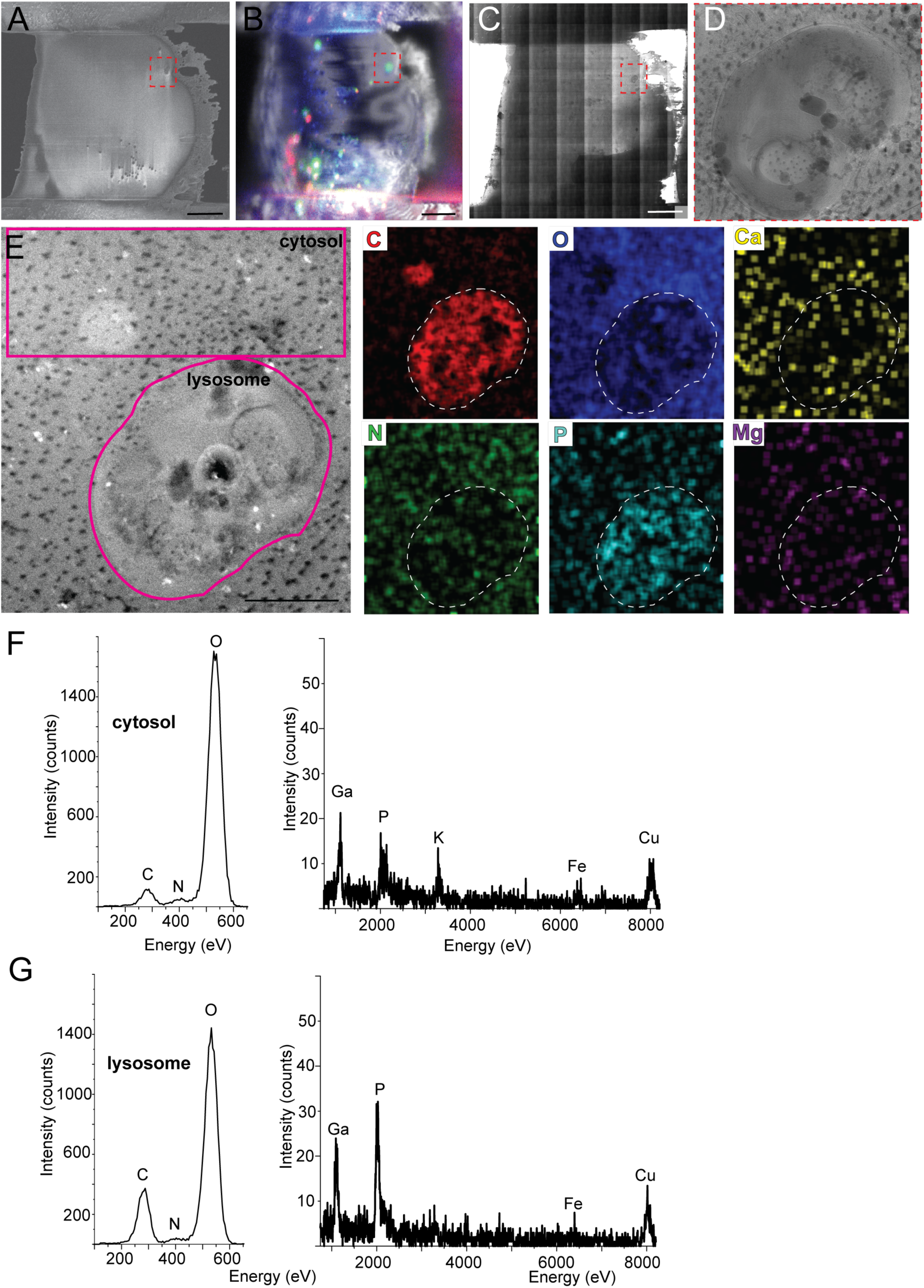
Energy-dispersive X-ray (EDX) spectroscopy combined with STEM imaging of a lysosome. (A) Scanning electron microscopy (SEM) images of lamella with visible droplets in inset. (B) SEM image superimposed with fluorescent image of green autofluorescence. (C) Overview image of corresponding recorded transmission electron microscope (TEM). (D) Large vesicular lysosome structure in higher magnification TEM image. (E) HAADF-STEM image of lysosome corresponding elemental images of carbon (C), oxygen (O), calcium (Ca), nitrogen (N), phosphorus (P) and magnesium (Mg). Scale bar 500 nm. The image shows two insets of the cytosol and droplet region used for subsequent averaging. EDX spectra of (F) cytosol region and (G) lysosome regions show significant gallium signal originating from the FIB preparation procedure.

**Supplementary Figure 9:**
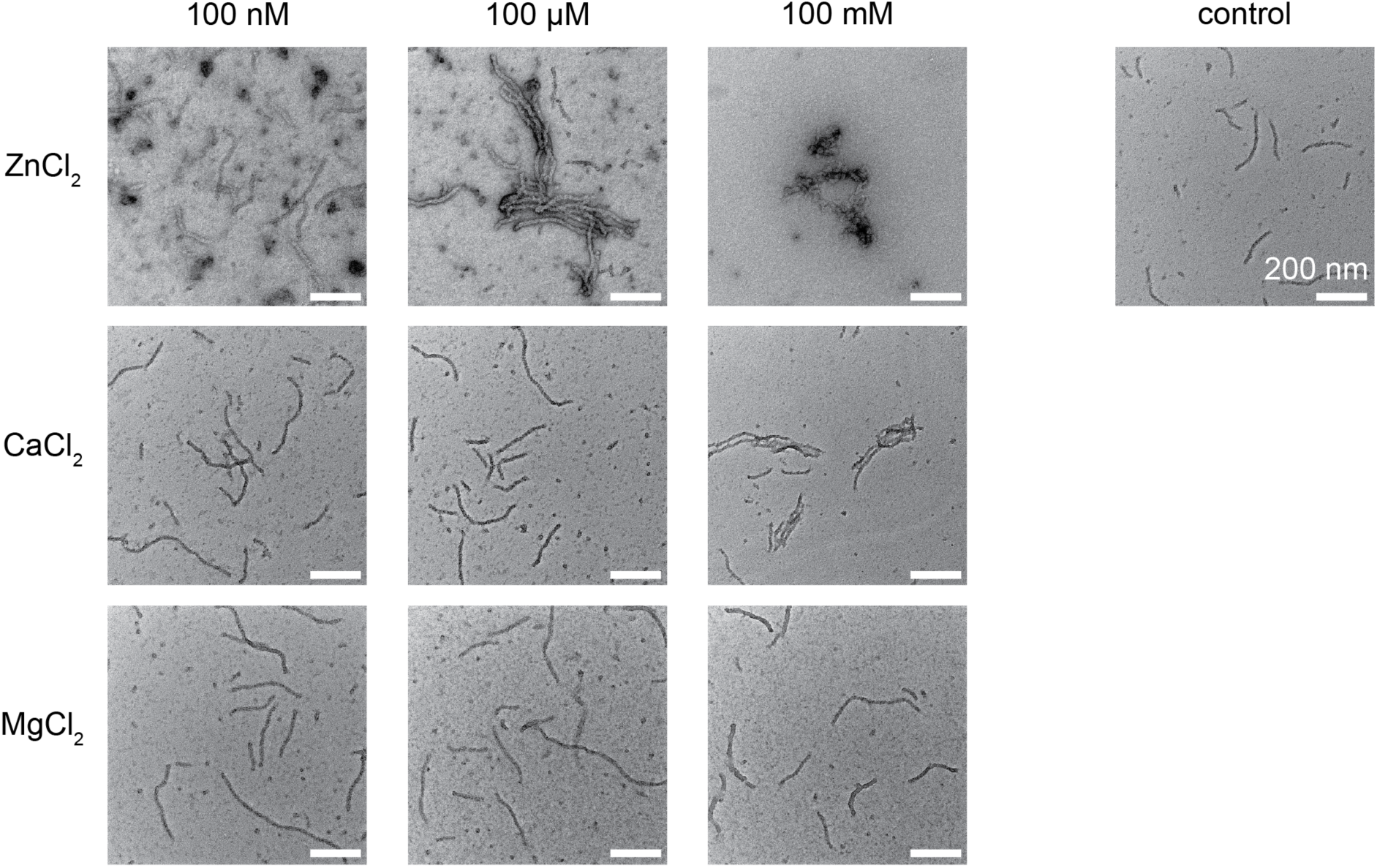
Effect of divalent cation salts on p62 filaments. p62 filaments at 1 µM concentration were mixed with 100 nM, 100 µM and 100 mM of ZnCl_2_, CaCl_2_ and MgCl_2_, respectively. The samples were incubated for 30 minutes and subsequently negatively stained. As a control, p62 filaments were mixed with desalting buffer. For increasing concentrations of ZnCl_2_from 100 µM and for CaCl_2_ from 100 mM bundling of p62 filaments was observed. For MgCl_2_ no clear effect of bundling was observed.

## Materials & Methods

### Protein expression, purification of p62, 4x-linear-ubiquitin and LC3B

Full-length p62 (440 residues) was cloned into a pETM43 expression vector containing an N-terminal maltose-binding-protein (MBP) tag and a C-terminal His-tag. Both tags featured a recognition sequence for HRV-3C protease, facilitating their removal after purification. 4x-linear-Ubiquitin, CFP-LC3B, and LC3B with an additional C-terminal cysteine residue were cloned into pGEX expression vector containing an N-terminal Glutathione-S-Transferase (GST) tag with HRV-3C protease cleavage site. For protein expression E. *coli* BL21 (DE3) cells were used, with bacterial cultures grown at 37°C in Terrific Broth (TB) medium to an OD_600_ of ∼2. Induction with IsoPropylThioGalactoside (IPTG) followed and cells were left to grow overnight at 23°C. Cells were harvested and the pellet was stored at −80 °C until further use.

After thawing on ice, cells were resuspended in lysis buffer [50 mM HEPES, pH 8.0, 300 mM NaCl, 2.5 mM MgSO_4_, 10 μM ZnCl_2_, 0.5 mM TCEP and 1 tablet EDTA-free protease inhibitor cocktail] and lysed using a cell disruptor at 1.85 bar. Cell debris was removed by centrifugation and the cleared lysate was applied to a Ni-NTA affinity column. The column was washed twice with lysis buffer containing 10 mM and 20 mM imidazole, respectively, and protein was eluted with lysis buffer containing 300 mM imidazole. Eluted protein fractions were pooled and subsequently applied to an amylose resin that binds MBP-fusion proteins with high affinity. The column was washed with lysis buffer and the protein was eluted with lysis buffer containing 25 mM maltose. Eluted protein fractions were pooled, and buffer was exchanged to desalting buffer [50 mM HEPES, pH 8.0, 300 mM NaCl and 0.5 mM TCEP] using Zeba desalting columns.

For the purification of LC3b and GST-4xUb, the cell pellets were resuspended in lysis buffer [50 mM HEPES, pH 7.4, 300 mM NaCl, 1 mM DTT, and 1 tablet EDTA-free protease inhibitor cocktail] and lysed using a cell disruptor at 1.9 bar. The cleared lysate was applied to Glutathione Sepharose 4 Fast Flow GST-tagged protein purification resin (Cytiva, Cat No 17513201), washed with 10 column volume lysis buffer, and eluted in lysis buffer containing 10 mM reduced glutathione. Elution fractions containing the protein of interested were pooled, further purified by size exclusion chromatography via SD200 16/600 column (Cytiva) [in 50 mM HEPES, pH 7.4, 150 mM NaCl, 1 mM DTT], and frozen at −80°C for storage.

### p62 filament formation and labeling

Protein was concentrated to approximately 2 mg/mL using a 30 kDa Amicon concentrator and subsequently incubated with 1:2.5 mass ratio of GST-HRV-3C protease for 3 h at room temperature and overnight at 4°C to remove the MBP and His-tag, allowing p62 filament formation. Filaments were isolated by centrifugation at 20,000 x g for 30 min and resuspended in desalting buffer. Isolated filaments were labeled to lysine residues by incubating with 1:1 molar ratio of Cy5-NHS for 1 h at room temperature and overnight at 4°C. Excess dye was removed by centrifugation, labeled filaments were resuspended in storage buffer [50 mM HEPES, pH 8.0, 50 mM NaCl and 0.5 mM TCEP] and stored at 4°C until further use.

### Cryo-EM structure determination of p62 filaments

Quantifoil Cu 200 mesh R1.2/R1.3 were glow discharged, and 2-5μL of the protein sample was applied in a Vitrobot mark IV (ThermoFischer Scientific). The sample chamber was maintained at 70-100 % humidity and 4-10 °C, with a 2-6 s blotting time before plunging into ethane/propane mixture. Data collection was performed using SerialEM(Mastronarde, 2005) on a Talos Arctica 200 KeV G2 transmission electron microscope (ThermoFischer Scientific), equipped with a K3 direct electron detector camera and Quantum GIF filter (Gatan, Inc). Approximately 4000 micrographs were collected in counted superresolution mode at 100,000x magnification with an underfocus range of −0.5 μm to −3 μm. The micrographs were motion corrected in WARP(Tegunov & Cramer, 2019) and particles were picked using the deep-learning based object detection software crYOLO (Wagner et al., 2019). Initially, the picked particle locations and micrographs were imported into RELION and SPRING(Desfosses et al., 2014; Scheres, 2012) that used CTFFIND4 for CTF estimation(Rohou & Grigorieff, 2015). Later, CTF corrected micrographs were imported in cryoSPARC (Punjani et al., 2017) and particle locations were extracted with a 90 % overlap yielding approximately 1,000,000 segments of a box size of 503 Å. Two rounds of 2D classification were performed in order to remove heterogeneous particles. Classes with clearly defined PB1 domains were then selected, leading to a total of 600,000 segments. The helical parameters as determined by Jakobi et al. (Jakobi et al., 2020) of truncated p62 helical filaments were taken as starting conditions for helical refinement in cryoSPARC, where the symmetry parameters converged. The final resolution was estimated by FSC using the 0.143 cutoff(Rosenthal & Henderson, 2003) and FDR-FSC criterion(Beckers & Sachse, 2020). Maps were rendered in UCSF ChimeraX (Pettersen et al., 2004).

### Single-particle fluorescence imaging in TIRF mode

Surface imaging was performed with an inverted microscope (Olympus IX-81, Olympus Corp., Shinjuku, Japan) in total internal reflection fluorescence (TIRF) mode. Molecules were excited using lasers at 488 and 639 nm (Sapphire 488-200 and Obis 637-140, Coherent Inc.) focused off-axis into the back-focal plane of an oil-immersion objective (Olympus UApoN, 100x, NA=1.49, Olympus Corp). The excitation power and corresponding intensity at the sample stage were 3.25 mW and 10.2 W/cm^2^ for the blue laser, and 2.2 mW and 6.8 W/cm^2^ for the red laser. The imaging was performed on an EMCCD camera (Andor iXon Ultra 888, Andor Technology) with 50, 100, 40 and 100 ms exposure time for Supplementary Videos 1-4, corresponding to 20, 10, 25 and 10 fps. Midway through acquiring Supplementary Video 2, the illumination was switched from blue to red laser to observe the same p62 filament with LC3b bound to it. Each video consists of 200 frames. Image analysis and video processing were carried out using Fiji (ImageJ, version 2.14.0) (Schindelin et al., 2012). The emitted fluorescence was collected with the same objective (objective based TIRF) and passed through a dual-band dichroic mirror as well as a dual-band emission filter (ZT488/640rpc and ZET488/640m, Chroma Technology Corp). p62 filaments-Cy5 and CFP-LC3b were diluted to concentrations of a few nM, and mixed in a 1:1 molar ratio. GST-4xUb was added at a concentration of 5 µM.

### Negative staining electron microscopy

For negative staining, 3.5 µL of the sample was applied onto glow-discharged carbon-formvar-coated 300-mesh copper grids and allowed to absorb for 2 min before blotted according to the side-blotting method. The sample was subsequently stained for 30 s with 6 µL 2% uranyl acetate, blotted and allowed to dry. Negatively stained samples were examined on a Talos L120C G2 transmission electron microscope (ThermoFisher Scientific), operated at 120 kV. Micrographs at high magnification (45,000x and 92,000x) were collected on a 4k x 4k Ceta 16M CEMOS camera using TEM Imaging & Analysis Software (TIA, ThermoFisher Scientific).

### Cryo-electron tomography of reconstituted p62 filaments

p62 filament cryo-EM grids were prepared as described above. Tilt series were recorded on a 200 kV Talos Arctica G2 transmission electron microscope (ThermoFisher Scientific), equipped with a Bioquantum GIF (Gatan, Inc) a Volta Phase Plate and a K3 direct electron detector (Gatan, Inc) using SerialEM (Mastronarde, 2005). Each tilt was recorded as movie at 63.000x magnification with a 0.681 Å/pixel size in super resolution mode and a dose of approximately 2.0 e/Å^2^/tilt, 10 frames per tilt, and a total dose per tilt series of approximately 130 e/Å^2^. All tilt series were recorded from −54° to 54° with 3° increments, using a dose-symmetric bidirectional acquisition scheme and a nominal defocus of −1.2 μm. The raw data was motion corrected and CTF corrected using WARP (version 1.0.9)(Tegunov & Cramer, 2019) and the tilt series were reconstructed using IMOD(Mastronarde & Held, 2017).

GST-4x-linear-Ubiquitin and p62 filaments were prepared as described above. 15 µM of GST-4x-linear-Ubiquitin were mixed with 1 µM p62 filaments. After 4 minutes incubation time at 4°C, the formed phase separation droplets were negatively stained as described above or plunge frozen with a Vitrobot Mark IV (ThermoFisher Scientific). 50 nm SUVs were prepared containing 5% DSPE-PEG(2000) maleimide (Avanti Polar lipids, Cat No 880126) in DOPS (Avantiv Polar lipids, Cat No 840035). The liposomes were incubated with 100 µM LC3b-Cys for 16 h at 4°C before removing unbound LC3b-Cys by size exclusion chromatography on Sepharcyl S-500 at Äkta Micro (Cytiva). A total of 2 mg/ml of liposomes were mixed with 1 µM p62 filaments and incubated for 5 min at 4 °C. The sample was plunge frozen at a Leica GM2 plunge freezer (Leica). Tomograms were collected at a Titan Krios 300 keV TEM equipped with an BioContinuum energy filter and K3 camera at 64,000x using Tomo5 software (ThermoFisher Scientific). The tilt series were recorded from −60° to 60° with a 3° increment in a dose-symmetric bidirectional acquisition scheme. An electron dose of 4.0 e^-^/Å^2^/tilt with 6 frames per tilt, resulted in a total dose per tilt series of approximately 160 e^-^/Å^2^. The defocus was varied between −2.0 to −4.0 μm. The images were motion corrected in WARP (Tegunov & Cramer, 2019) and tomograms reconstructed with AreTomo (S. Zheng et al., 2022). Overall binning was 16x and resulted in a pixel size of 11.2 Å. Tomograms were denoised and missing wedge corrected with IsoNet (Liu et al., 2021).

### Cell culture and transfection

Cell culture experiments were carried out in human RPE1 cells (hTERT RPE1, CRL-4000, ATCC) cultured at 37°C and 5% CO_2_ in DMEM GlutaMAX media (Gibco, Cat No 31331028) supplemented with 10% FCS (Sigma-Aldrich, Cat No F7524) and Pen/Strep (Gibco, Cat No 15140122). For confocal microscopy experiments prior to plunge-freezing, the culturing media was substituted with media of low fluorescence background; Fluorobrite DMEM (Gibco, Cat No A1896701). Additional experiments were performed on RPE1 cells stably expressing human mCherry-p62. Transient expression of mCherry-p62 was executed by transfection with FugeneHD (Promega, Cat No E2311) and a modified pDEST vector containing the human wildtype p62 gene fused with an N-terminal mCherry sequence (Pankiv et al., 2007). Knockdown experiments were carried out by transfection with RNAiMAX (ThermoFisher, Cat No 13778075) and ON-TARGETplus SMARTpool siRNA against ATG5 (Dharmcon Horizon Discovery, Cat No L-004374-00-0005), GADPH (Dharmcon Horizon Discovery, Cat No D-001830-10-05) and a non-targeting control (Dharmcon Horizon Discovery, Cat No D-001810-10-05). Follow-up experiments were performed 72 hours after a reverse transfection with 75 nm siRNA. Lipid droplets were stained with 1:500 Lipi-Blue (Dojindo, Cat No LD01-10) for 24 hours. Lysosomes were stained with the CytoPainter Lysosomal staining kit (Abcam, Cat No ab112136).

### Western Blot

Cells were lysed by a quick wash with ice cold PBS (Gibco, Cat No 10010056) and a subsequent 15 min incubation in ice cold RIPA buffer (Sigma-Aldrich, Cat No R0278-50mL) supplemented with a protease inhibitor cocktail (Roche, Cat No 11836170001). Cell lysate was collected with a cell scraper before removing the cell debris by centrifugation. Total protein content was quantified and corrected using a Pierce 660 nm assay (ThermoFisher, Cat No 22662) before boiling the sampling in a reducing sample buffer and separation on a 4-15 % mini-PROTEAN Precast SDS-PAGE (Bio-Rad, Cat No 4561085). Transfer to PVDF membrane was performed using a Trans-Blot pack (Bio-Rad, Cat No 1704156) and the membrane was blocked for 2 h in 2 % BSA in TBST at 4 °C. The membrane was incubated overnight with primary antibody in 1 % BSA in TBST at 4 °C, washed 3 times in TBST before the secondary antibody was added for 2 h in 1 % BSA in TBST at 4 °C. Prior to imaging on the Azure Sapphire RGB Biomolecular Imager (Azure Biosystems), the membrane was washed 3 times in TBST and incubated in Azure Radiance Q HRP (Biozym, Cat No 512101). The following antibodies were used: polyclonal rabbit anti-p62 (MBL International, Cat No PM045), monoclonal rabbit anti-ATG5 (Abcam, Cat No ab108327), monoclonal mouse anti-GADPH (Invitrogen, Cat No MA1-16757), polyclonal rabbit anti-LC3 (Abcam, Cat No ab48394), goat anti-mouse HRP conjugate (ThermoFisher Cat No 32230), and goat anti-rabbit HRP conjugate (ThermoFisher, Cat No 31460). Band intensity quantifications were performed using ImageLab (BioRad, version 6.1.0).

### Cellular tomography grid preparation

RPE1 Cells were seeded on micropatterned grids before plunge freezing, For a more detailed protocol see Berkamp et al. Bio-Protocols 2023 (Berkamp et al., 2023; Toro-Nahuelpan et al., 2019). Briefly, 200 mesh gold Quantifoil R2/1 grids with a SiO_2_ support layer (Quantifoil, Cat No N1-S15nAu20-01) were glow discharged and treated with PLL-g-PEG (SuSoS, PLL(20)-g[3.5]-PEG(5)) overnight. Next, the grids were washed in PBS and placed in drops of 14.5 mg/mL PLPP (Alvéole, Cat No PLPP) and 20 μm circles in the center of 10x10 grid squares were micropatterned with a PRIMO photopatterning device (Alvéole, Photopatterning Device consisting of a DMD + laser 375 nm, 75 mW module, and driving electronics). The grids were then washed thoroughly in PBS and incubated in human fibronectin immediately before use (Gibco, Cat No PHE0023). The cells were trypsinized (Gibco, Cat No 25300054), passed through a 40 μm cell strainer (PluriSelect, Cat No 43-10040-40) and counted before seeding ∼200,000 cells in a 30 mm glass-bottom dish (Greiner Bio-One, Cat No 627860) containing the micropatterned grids in a small amount of DMEM. The cells were allowed to attach to the grids for 4-5 h before vitrification in liquid ethane using a Vitrobot Mark IV (ThermoFisher Scientific). The following settings were used; a chamber temperature of 37 °C, a chamber humidity of 85 % humidity, backside blotting with a 8.5 s blot time and blot force of −10 and a drain time of 1 s.

### Confocal microscopy

RPE1 cell phenotypes were studied using a Zeiss LSM710 laser scanning confocal microscope (Carl Zeiss GmbH). The cells were grown in 30 mm glass bottom dishes with 4 compartments (Greiner Bio-One, Cat No 627870) and imaged at 37°C using a Zeiss 63× oil immersion objective (Plan-Apochromat, DIC, 1.4 NA). Lipi-Blue was excited with a diode laser at 405 nm and fluorescent emission was detected after passing a suitable emission filter (410-500 nm) using a Multi Alkali photomultiplier tube. The lysosomal stain was excited using an argon ion laser at 488 nm and emission was detected using an emission filter (499-541 nm). MCherry-p62 was detected using a 543 HeNe543 laser and an emission filter set to 579-696 nm using a GaAsP photomultiplier tube. Image analysis and quantification were done using Fiji (ImageJ, version 2.14.0) (Schindelin et al., 2012) and a self-written routine in CellProfiler (version 4.2.6) (Stirling et al., 2021). Measurements were exported as .csv files and analyzed in OriginPro 2023 (OriginLab Corporation, version 10.0.0.154). Vitrified grids and thinned lamellae were studied using a Zeiss LSM800 upright laser scanning confocal microscope with a cryostage (Linkam CMS196). Super resolution images were acquired using a Zeiss 100x air immersion objective (EC Epiplan NeoFluar, DIC, NA=0.75) and an AiryScan Detector. MCherry-p62 was detected using a 561nm diode. Image processing was done using Zen Blue (Carl Zeiss GmbH, version 3.3.89).

### Correlative light and electron microscopy

Cryo-lamellae were made in a correlative manner using the in-chamber METEOR fluorescence microscope (Delmic, Cat No 2707-999-0014-2) equipped with a LMPLFLN 50×, NA = 0.5, WD = 10.6 mm objective lens in the Aquilos2 cryo-FIB-SEM system (ThermoFisher Scientific). In short; micropatterned grids with RPE1 cells were loaded under cryogenic conditions taking special care to avoid contamination build-up from air humidity. Cells were located using the electron beam and the sample was placed at eucentric height using MAPS software (ThermoFisher Scientific, version 3.19). Cell phenotype was confirmed by imaging each cell in the METEOR fluorescence microscope. Each cell was imaged using the focused ion beam and the location of the final 150 nm lamella was marked in AutoTEM (ThermoFisher Scientific, version 2.4.1). The lamellae were generated using a step-wise process; the cell was thinned to 1 μm with a 0.5 nA milling current, then thinned to 600 nm with a 0.3 nA current, next to 300 nm with a 0.1 nA current and finally thinned to 200 nm with a 50 pA ion beam. The final polishing step in which the lamella was thinned to 120-180 nm took place with a 30 pA ion beam. The final lamella was once again imaged using the METEOR fluorescence microscope and these images were used to determine where tomograms would be recorded. The lamella were coated with a thin platinum layer; sputtered for 7 seconds at 7 mA, to prevent sample charging. For a more detailed protocol see Berkamp et al. Bio-Protocols 2023 (Berkamp et al., 2023). Next, the samples were transferred under cryo-conditions to a 200 kV Talos Arctica G2 transmission electron microscope (ThermoFisher Scientific), equipped with a Bioquantum GIF (Gatan, Inc) and a K3 direct electron detector (Gatan, Inc). Tilt series were recorded on p62 positive areas of the lamellae, using SerialEM (version 4.2.0)(Mastronarde, 2005) as movies at 24,000x magnification with a 1.7435 Å/pixel size in super resolution mode and a dose of approximately 2.2 e/Å^2^/tilt, 10 frames per tilt, and a total dose per tilt series of approximately 130 e/Å^2^. All tilt series were recorded from −54° to 54° with 3° increments starting at the pre-tilt of the lamella, using a dose-symmetric bidirectional acquisition scheme with weighted dose according to tilt angle and a nominal defocus of −7 μm. The raw data was motion corrected and CTF corrected using WARP (version 1.0.9) (Tegunov & Cramer, 2019) and the tilt series were reconstructed using Aretomo2 (SBGrid) (Morin et al., 2013; S. Zheng et al., 2022). To aid the cryo tomogram visualization, they were deconvolved, denoised and the missing wedge compensated using IsoNet (Liu et al., 2021) and cryoCARE (Buchholz et al., 2019). To aid in segmentation a pre-trained U-Net model for non-denoised data in MemBrain-Seg (version 9b) (Lamm et al., 2022) was used and the processed tomograms were then loaded into Amira 3D (ThermoFisher Scientific, version 2022.2) for semi-manual segmentation. Tomograms were visualized using the IMOD software package (version 4.11.20) (Mastronarde & Held, 2017). To aid in targeting the EDX spectra, high dose, single tilt micrographs were recorded on p62 positive areas of the lamellae at 24,000x and a dose of approximately 40 e^-^/Å^2^/micrograph and 180 frames per tilt with each a dose of approximately 0.24 e^-^/Å^2^. Motion correction for these micrographs was done in Relion (SBGrid, version 4.0.1) (Kimanius et al., 2021)

### Energy-dispersive X-ray (EDX) spectroscopy

The in situ EDX experiments were carried out on a probe-corrected 200 kV FEI Titan microscope (ThermoFisher Scientific) fitted with a Super-X G2 quadrant detector EDX system (Bruker) and a Simple Origin dual grid cryo transfer holder (Simple Origin, Inc). The microscope was operated under cryo-conditions in STEM mode with a 1.5 mrad convergence angle, a 1 nm probe size at a magnification of 55,000, resulting in a 2.9 nm pixel size. EDX spectra were recorded with a 50 μs dwell time and 50 frames resulting in a total dose of 500 e/Å^2^, a camera length of 3.5m and a detector dead time of approximately 70%. EDX spectra processing was done in Velox (ThermoFisher Scientific, version 3.14). Atomic map representations were rendered with a 12 px pre-filter average and a 12 px post-filter average for the p62 structures and a 25 px pre-filter average and a 9 px post-filter average for the lipid droplet and lysosome control experiments. The spectra were generated by segmenting the p62 coat in the HAADF-STEM micrograph and segmenting the same number of pixels in the lipid droplet core and a random area of the cytosol. This same process was followed for the control experiments.

## Acknowledgements

S.B. was supported by a fellowship from the Alexander von Humboldt Foundation. This work was supported by the European Research Council under Grant Agreement 101118656 (ERC Synergy project 4D-BioSTEM). The work was initially supported by the Boehringer Ingelheim Fonds Exploration grant. The authors would like to thank Andreas Brech and Sebastian Schultz from Oslo University Hospital, Oslo, Norway for providing the stable mCherry-p62 RPE1 cell line. LT acknowledges the support India Alliance DBT/Wellcome trust grant (IA/21/2/505925). The authors gratefully acknowledge the electron microscopy access time and computing time granted by the biological EM facility of the Ernst-Ruska Centre at Forschungszentrum Jülich. In particular, we thank Julio Ortiz for the recording of the Volta phase plate p62 filament tomograms. The authors gratefully acknowledge the computing time granted by the JARA Vergabegremium and provided on the JARA Partition part of the supercomputer JURECA at Forschungszentrum Jülich (Thörnig, 2021).

## Author Contributions

SB, SM, LJ and CS designed the project. SM, AK, LJ purified p62 filaments. SM and AK determined the cryo-EM structure of p62. LJ and AK reconstituted p62 filaments with binding partners and LC3b conjugated liposomes. AK, OK and JF performed single-particle fluorescence measurements and analysis. SB performed the live cell confocal imaging and data analysis. SB performed in situ CLEM followed by cryo-ET analysis. SB, PL and RDB performed the EDX measurements and analus. SB, LJ, AK and CS wrote the manuscript with input from the other authors.

## Data and code availability

The EMDB accession number for the p62 cryo-EM map is EMD-52134 and the corresponding PDB-ID 9HGE for the fitted PB1 coordinates.

